## Supplementary material for "Prophage regulation of *Shewanella fidelis* 3313 motility and biofilm formation: implications for gut colonization dynamics in *Ciona robusta*": Tex: elife-sample.bib

@article{RN102,
   author = {Abusleme, L and Moutsopoulos, NM},
   title = {IL-17: overview and role in oral immunity and microbiome},
   journal = {Oral Diseases},
   volume = {23},
   number = {7},
   pages = {854-865},
   ISSN = {1354-523X},
   DOI = {https://doi.org/10.1111/odi.12598},
   url = {https://onlinelibrary.wiley.com/doi/abs/10.1111/odi.12598},
   year = {2017},
   type = {Journal Article}
}

@article{RN55,
   author = {Ackermann, Hans-W and Heldal, Mikal},
   title = {Basic electron microscopy of aquatic viruses},
   journal = {Manual of aquatic viral ecology},
   volume = {18},
   pages = {182-192},
   year = {2010},
   type = {Journal Article}
}

@article{RN94,
   author = {Akhter, Sajia and Aziz, Ramy K. and Edwards, Robert A.},
   title = {PhiSpy: a novel algorithm for finding prophages in bacterial genomes that combines similarity- and composition-based strategies},
   journal = {Nucleic Acids Research},
   volume = {40},
   number = {16},
   pages = {e126-e126},
   note = {OA status: green_published},
   ISSN = {1362-4962},
   DOI = {10.1093/nar/gks406},
   url = {http://europepmc.org/articles/pmc3439882?pdf=render},
   year = {2012},
   type = {Journal Article}
}

@article{RN18,
   author = {Allen, H. K. and Looft, T. and Bayles, D. O. and Humphrey, S. and Levine, U. Y. and Alt, D. and Stanton, T. B.},
   title = {Antibiotics in feed induce prophages in swine fecal microbiomes},
   journal = {MBio},
   url = {http://www.ncbi.nlm.nih.gov/pubmed/22128350},
   year = {2011}
}

@article{RN99,
   author = {Andersen, Claus Lindbjerg and Jensen, Jens Ledet and Ørntoft, Torben Falck},
   title = {Normalization of real-time quantitative reverse transcription-PCR data: a model-based variance estimation approach to identify genes suited for normalization, applied to bladder and colon cancer data sets},
   journal = {Cancer research},
   volume = {64},
   number = {15},
   pages = {5245-5250},
   ISSN = {0008-5472},
   year = {2004}
}

@article{RN50,
   author = {Arndt, David and Grant, Jason R. and Marcu, Ana and Sajed, Tanvir and Pon, Allison and Liang, Yongjie and Wishart, David S.},
   title = {PHASTER: a better, faster version of the PHAST phage search tool},
   journal = {Nucleic Acids Research},
   volume = {44},
   number = {W1},
   pages = {W16-W21},
   ISSN = {0305-1048},
   DOI = {10.1093/nar/gkw387},
   url = {https://dx.doi.org/10.1093/nar/gkw387},
   year = {2016},
   type = {Journal Article}
}

@article{RN93,
   author = {Aziz, Ramy K and Bartels, Daniela and Best, Aaron A and Dejongh, Matthew and Disz, Terrence and Edwards, Robert A and Formsma, Kevin and Gerdes, Svetlana and Glass, Elizabeth M and Kubal, Michael and Meyer, Folker and Olsen, Gary J and Olson, Robert and Osterman, Andrei L and Overbeek, Ross A and Mcneil, Leslie K and Paarmann, Daniel and Paczian, Tobias and Parrello, Bruce and Pusch, Gordon D and Reich, Claudia and Stevens, Rick and Vassieva, Olga and Vonstein, Veronika and Wilke, Andreas and Zagnitko, Olga},
   title = {The RAST Server: Rapid Annotations using Subsystems Technology},
   journal = {BMC Genomics},
   volume = {9},
   number = {1},
   pages = {75},
   note = {OA status: gold_doaj},
   ISSN = {1471-2164},
   DOI = {10.1186/1471-2164-9-75},
   url = {https://doi.org/10.1186/1471-2164-9-75},
   year = {2008},
   type = {Journal Article}
}

@article{RN95,
   author = {Aziz, Ramy K and Edwards, Robert A and Taylor, William W and Low, Donald E and McGeer, Allison and Kotb, Malak},
   title = {Mosaic prophages with horizontally acquired genes account for the emergence and diversification of the globally disseminated M1T1 clone of Streptococcus pyogenes},
   journal = {Journal of bacteriology},
   volume = {187},
   number = {10},
   pages = {3311-3318},
   ISSN = {0021-9193},
   year = {2005}
}

@article{RN19,
   author = {Banks, D. J. and Lei, B. and Musser, J. M.},
   title = {Prophage induction and expression of prophage-encoded virulence factors in group A \textit{Streptococcus serotype} M3 strain MGAS315},
   journal = {Infect Immun},
   volume = {71},
   number = {12},
   pages = {7079-86},
   ISSN = {0019-9567 (Print)
0019-9567 (Linking)},
   url = {http://www.ncbi.nlm.nih.gov/pubmed/14638798},
   year = {2003}
}

@article{RN101,
   author = {Barbier, Mariette and Damron, F. Heath},
   title = {Rainbow Vectors for Broad-Range Bacterial Fluorescence Labeling},
   journal = {PLOS ONE},
   volume = {11},
   number = {3},
   pages = {e0146827},
   ISSN = {1932-6203},
   DOI = {10.1371/journal.pone.0146827},
   url = {https://dx.doi.org/10.1371/journal.pone.0146827},
   year = {2016},
   type = {Journal Article}
}

@article{RN104,
   author = {Barbier, Mariette and Damron, F Heath},
   title = {Rainbow vectors for broad-range bacterial fluorescence labeling},
   journal = {PLoS One},
   volume = {11},
   number = {3},
   pages = {e0146827},
   ISSN = {1932-6203},
   year = {2016},
   type = {Journal Article}
}

@article{RN89,
   author = {Barr, J. J. and Auro, R. and Furlan, M. and Whiteson, K. L. and Erb, M. L. and Pogliano, J. and Stotland, A. and Wolkowicz, R. and Cutting, A. S. and Doran, K. S. and Salamon, P. and Youle, M. and Rohwer, F.},
   title = {Bacteriophage adhering to mucus provide a non-host-derived immunity},
   journal = {Proc Natl Acad Sci U S A},
   volume = {110},
   number = {26},
   pages = {10771-6},
   ISSN = {1091-6490 (Electronic)
0027-8424 (Linking)},
   DOI = {10.1073/pnas.1305923110},
   url = {http://www.ncbi.nlm.nih.gov/pubmed/23690590},
   year = {2013}
}

@article{RN44,
   author = {Bolger, A. M. and Lohse, M. and Usadel, B.},
   title = {Trimmomatic: a flexible trimmer for Illumina sequence data},
   journal = {Bioinformatics},
   volume = {30},
   number = {15},
   pages = {2114-20},
   ISSN = {1367-4811 (Electronic)
1367-4803 (Linking)},
   DOI = {10.1093/bioinformatics/btu170},
   url = {https://www.ncbi.nlm.nih.gov/pubmed/24695404},
   year = {2014},
   type = {Journal Article}
}

@article{RN10,
   author = {Boling, L. and Cuevas, D. A. and Grasis, J. A. and Kang, H. S. and Knowles, B. and Levi, K. and Maughan, H. and McNair, K. and Rojas, M. I. and Sanchez, S. E. and Smurthwaite, C. and Rohwer, F.},
   title = {Dietary prophage inducers and antimicrobials: toward landscaping the human gut microbiome},
   journal = {Gut Microbes},
   volume = {11},
   number = {4},
   pages = {721-734},
   ISSN = {1949-0984 (Electronic)
1949-0976 (Linking)},
   DOI = {10.1080/19490976.2019.1701353},
   url = {https://www.ncbi.nlm.nih.gov/pubmed/31931655},
   year = {2020}
}

@article{RN80,
   author = {Bollinger, R. R. and Everett, M. L. and Wahl, S. D. and Lee, Y. H. and Orndorff, P. E. and Parker, W.},
   title = {Secretory IgA and mucin-mediated biofilm formation by environmental strains of \textit{Escherichia coli}: role of type 1 pili},
   journal = {Mol Immunol},
   volume = {43},
   number = {4},
   pages = {378-87},
   ISSN = {0161-5890 (Print)
0161-5890 (Linking)},
   DOI = {10.1016/j.molimm.2005.02.013},
   url = {http://www.ncbi.nlm.nih.gov/pubmed/16310051},
   year = {2006}
}

@article{RN12,
   author = {Bondy-Denomy, Joseph and Davidson, Alan R},
   title = {When a virus is not a parasite: the beneficial effects of prophages on bacterial fitness},
   journal = {Journal of microbiology},
   volume = {52},
   number = {3},
   pages = {235-242},
   ISSN = {1225-8873},
   year = {2014}
}

@article{RN63,
   author = {Boulos, L},
   title = {Pré vost M, Barbeau B, Coallier J, Desjardins R. LIVE/DEAD BacLight: application of a new rapid staining method for direct enumeration of viable and total bacteria in drinking water},
   journal = {J Microbiol Methods},
   volume = {37},
   pages = {77-86},
   year = {1999},
   type = {Journal Article}
}

@article{RN52,
   author = {Bryksin, Anton V. Matsumura, Ichiro},
   title = {Overlap extension PCR cloning: a simple and reliable way to create recombinant plasmids},
   journal = {BioTechniques},
   volume = {48},
   number = {6},
   pages = {463-465},
   ISSN = {0736-6205},
   DOI = {10.2144/000113418},
   url = {https://dx.doi.org/10.2144/000113418},
   year = {2010}
}

@article{RN73,
   author = {Chao, L. and Rakshe, S. and Leff, M. and Spormann, A. M.},
   title = {PdeB, a Cyclic Di-GMP-Specific Phosphodiesterase That Regulates \textit{Shewanella oneidensis} MR-1 Motility and Biofilm Formation},
   journal = {Journal of Bacteriology},
   volume = {195},
   number = {17},
   pages = {3827-3833},
   ISSN = {0021-9193},
   DOI = {10.1128/jb.00498-13},
   url = {https://dx.doi.org/10.1128/jb.00498-13},
   year = {2013}
}

@article{RN34,
   author = {Chiba, S. and Sasaki, A. and Nakayama, A. and Takamura, K. and Satoh, N.},
   title = {Development of \textit{Ciona intestinalis} juveniles (through 2nd ascidian stage)},
   journal = {Zoolog Sci},
   volume = {21},
   number = {3},
   pages = {285-98},
   ISSN = {0289-0003 (Print)
0289-0003 (Linking)},
   DOI = {10.2108/zsj.21.285},
   url = {https://www.ncbi.nlm.nih.gov/pubmed/15056923},
   year = {2004}
}

@misc{RN59,
   author = {Cirino, Paola and Toscano, Alfonso and Caramiello, Davide and Macina, Alberto and Miraglia, Valentina and Monte, Assunta},
   title = {Laboratory culture of the ascidian \textit{Ciona intestinalis}(L.): a model system for molecular developmental biology research},
   url = {http://www.mbl.edu/html/BB/MMER/CIR/CirTit.html [year, month, day]},
   publisher={University of Chicago, Marine Biological Laboratory},
   year = {2002}
}

@article{RN53,
   author = {Cochran, Pk and Kellogg, Ca and Paul, Jh},
   title = {Prophage induction of indigenous marine lysogenic bacteria by environmental pollutants},
   journal = {Marine Ecology Progress Series},
   volume = {164},
   pages = {125-133},
   ISSN = {0171-8630},
   DOI = {10.3354/meps164125},
   url = {https://dx.doi.org/10.3354/meps164125},
   year = {1998},
   type = {Journal Article}
}

@misc{RN70,
   author = {Constantin, Ahlmann-Eltze; Patil, Indrajeet},
   title = {{ggsignif}: R Package for Displaying Significance Brackets for {'ggplot2'}},
   DOI = {10.31234/osf.io/7awm6},
   url = {https://psyarxiv.com/7awm6},
    publisher={PsyArXiV},
   year = {2021}
}

@article{RN113,
   author = {Corts, Anna D and Thomason, Lynn C and Gill, Ryan T and Gralnick, Jeffrey A},
   title = {A new recombineering system for precise genome-editing in \textit{Shewanella oneidensis} strain MR-1 using single-stranded oligonucleotides},
   journal = {Scientific Reports},
   volume = {9},
   number = {1},
   pages = {39},
   ISSN = {2045-2322},
   year = {2019},
   type = {Journal Article}
}

@article{RN109,
   author = {Cua, Daniel J. and Tato, Cristina M.},
   title = {Innate IL-17-producing cells: the sentinels of the immune system},
   journal = {Nature Reviews Immunology},
   volume = {10},
   number = {7},
   pages = {479-489},
   ISSN = {1474-1733},
   DOI = {10.1038/nri2800},
   url = {https://dx.doi.org/10.1038/nri2800},
   year = {2010},
   type = {Journal Article}
}

@article{RN35,
   author = {Davidson, Brad},
   title = {\textit{Ciona intestinalis} as a model for cardiac development},
   journal = {Seminars in Cell \& Developmental Biology},
   volume = {18},
   number = {1},
   pages = {16-26},
   ISSN = {1084-9521},
   DOI = {10.1016/j.semcdb.2006.12.007},
   url = {https://dx.doi.org/10.1016/j.semcdb.2006.12.007},
   year = {2007}
}

@article{RN20,
   author = {Diard, M. and Bakkeren, E. and Cornuault, J. K. and Moor, K. and Hausmann, A. and Sellin, M. E. and Loverdo, C. and Aertsen, A. and Ackermann, M. and De Paepe, M. and Slack, E. and Hardt, W. D.},
   title = {Inflammation boosts bacteriophage transfer between \textit{Salmonella spp}},
   journal = {Science},
   volume = {355},
   number = {6330},
   pages = {1211-1215},
   ISSN = {1095-9203 (Electronic)
0036-8075 (Linking)},
   DOI = {10.1126/science.aaf8451},
   url = {http://www.ncbi.nlm.nih.gov/pubmed/28302859},
   year = {2017}
}

@article{RN84,
   author = {Dishaw, L. J. and Cannon, J. P. and Litman, G. W. and Parker, W.},
   title = {Immune-directed support of rich microbial communities in the gut has ancient roots},
   journal = {Dev Comp Immunol},
   ISSN = {1879-0089 (Electronic)
0145-305X (Linking)},
   DOI = {10.1016/j.dci.2014.06.011},
   url = {http://www.ncbi.nlm.nih.gov/pubmed/24984114},
   year = {2014}
}

@article{RN32,
   author = {Dishaw, L. J. and Flores-Torres, J. and Lax, S. and Gemayel, K. and Leigh, B. and Melillo, D. and Mueller, M. G. and Natale, L. and Zucchetti, I. and De Santis, R. and Pinto, M. R. and Litman, G. W. and Gilbert, J. A.},
   title = {The gut of geographically disparate \textit{Ciona intestinalis} harbors a core microbiota},
   journal = {PLoS One},
   volume = {9},
   number = {4},
   pages = {e93386},
   ISSN = {1932-6203 (Electronic)
1932-6203 (Linking)},
   DOI = {10.1371/journal.pone.0093386},
   url = {https://www.ncbi.nlm.nih.gov/pubmed/24695540},
   year = {2014}
}

@article{RN41,
   author = {Dishaw, Larry J. and Giacomelli, Stefano and Melillo, Daniela and Zucchetti, Ivana and Haire, Robert N. and Natale, Lenina and Russo, Nicola A. and De Santis, Rosaria and Litman, Gary W. and Pinto, Maria Rosaria},
   title = {A role for variable region-containing chitin-binding proteins (VCBPs) in host gut–bacteria interactions},
   journal = {Proceedings of the National Academy of Sciences},
   volume = {108},
   number = {40},
   pages = {16747-16752},
   DOI = {10.1073/pnas.1109687108},
   url = {https://www.pnas.org/content/pnas/108/40/16747.full.pdf},
   year = {2011}
}

@article{RN40,
   author = {Dishaw, L. J. and Leigh, B. and Cannon, J. P. and Liberti, A. and Mueller, M. G. and Skapura, D. P. and Karrer, C. R. and Pinto, M. R. and De Santis, R. and Litman, G. W.},
   title = {Gut immunity in a protochordate involves a secreted immunoglobulin-type mediator binding host chitin and bacteria},
   journal = {Nat Commun},
   volume = {7},
   pages = {10617},
   ISSN = {2041-1723 (Electronic)
2041-1723 (Linking)},
   DOI = {10.1038/ncomms10617},
   url = {http://www.ncbi.nlm.nih.gov/pubmed/26875669},
   year = {2016}
}

@article{RN83,
   author = {Donaldson, G. P. and Ladinsky, M. S. and Yu, K. B. and Sanders, J. G. and Yoo, B. B. and Chou, W. C. and Conner, M. E. and Earl, A. M. and Knight, R. and Bjorkman, P. J. and Mazmanian, S. K.},
   title = {Gut microbiota utilize immunoglobulin A for mucosal colonization},
   journal = {Science},
   ISSN = {1095-9203 (Electronic)
0036-8075 (Linking)},
   DOI = {10.1126/science.aaq0926},
   url = {http://www.ncbi.nlm.nih.gov/pubmed/29724905},
   year = {2018}
}

@misc{RN111,
   author = {Dotmatics},
   title = { SnapGene® software},
   publisher = {https://www.snapgene.com/},
   year = {2023},
   type = {Multimedia Application}
}

@article{RN3,
   author = {Duncan, K. and Carey-Ewend, K. and Vaishnava, S.},
   title = {Spatial analysis of gut microbiome reveals a distinct ecological niche associated with the mucus layer},
   journal = {Gut Microbes},
   volume = {13},
   number = {1},
   pages = {1874815},
   ISSN = {1949-0984 (Electronic)
1949-0976 (Linking)},
   DOI = {10.1080/19490976.2021.1874815},
   url = {https://www.ncbi.nlm.nih.gov/pubmed/33567985},
   year = {2021}
}

@misc{RN67,
   author = {Eklund, Aron; Trimble, James},
   title = {The Bee Swarm Plot, an Alternative to Stripchart},
   url = {https://github.com/aroneklund/beeswarm},
   year = {2021}
}

@article{RN21,
   author = {Fang, Y. and Mercer, R. G. and McMullen, L. M. and Ganzle, M. G.},
   title = {Induction of Shiga Toxin-Encoding Prophage by Abiotic Environmental Stress in Food},
   journal = {Appl Environ Microbiol},
   volume = {83},
   number = {19},
   DOI = {10.1128/AEM.01378-17},
   url = {http://www.ncbi.nlm.nih.gov/pubmed/28778890},
   year = {2017}
}

@article{RN27,
   author = {Fortier, L. C. and Sekulovic, O.},
   title = {Importance of prophages to evolution and virulence of bacterial pathogens},
   journal = {Virulence},
   volume = {4},
   number = {5},
   pages = {354-65},
   ISSN = {2150-5608 (Electronic)
2150-5594 (Linking)},
   DOI = {10.4161/viru.24498},
   url = {http://www.ncbi.nlm.nih.gov/pubmed/23611873},
   year = {2013}
}

@article{RN22,
   author = {Garcia-Russell, N. and Elrod, B. and Dominguez, K.},
   title = {Stress-induced prophage DNA replication in \textit{Salmonella enterica} serovar Typhimurium},
   journal = {Infect Genet Evol},
   volume = {9},
   number = {5},
   pages = {889-95},
   ISSN = {1567-7257 (Electronic)
1567-1348 (Linking)},
   DOI = {10.1016/j.meegid.2009.05.017},
   url = {http://www.ncbi.nlm.nih.gov/pubmed/19501196},
   year = {2009}
}

@article{RN61,
   author = {Gaudy Jr, AF and Abu-Niaaj, F and Gaudy, ET},
   title = {Statistical study of the spot-plate technique for viable-cell counts},
   journal = {Applied microbiology},
   volume = {11},
   number = {4},
   pages = {305-309},
   ISSN = {0003-6919},
   year = {1963}
}

@article{RN42,
   author = {Gödeke, Julia and Paul, Kristina and Lassak, Jürgen and Thormann, Kai M.},
   title = {Phage-induced lysis enhances biofilm formation in \textit{Shewanella oneidensis} MR-1},
   journal = {The ISME Journal},
   volume = {5},
   number = {4},
   pages = {613-626},
   ISSN = {1751-7370},
   DOI = {10.1038/ismej.2010.153},
   url = {https://doi.org/10.1038/ismej.2010.153 },
   year = {2011},
}

@article{RN4,
   author = {Harrington, V. and Lau, L. and Seddu, K. and Suez, J.},
   title = {Ecology and Medicine Converge at the Microbiome-Host Interface},
   journal = {mSystems},
   pages = {e0075621},
   ISSN = {2379-5077 (Print)
2379-5077 (Linking)},
   DOI = {10.1128/mSystems.00756-21},
   url = {https://www.ncbi.nlm.nih.gov/pubmed/34463567},
   year = {2021}
}

@inbook{RN71,
   author = {Harshey, Rasika M.},
   title = {Phage Mu},
   publisher = {Springer US},
   pages = {193-234},
   DOI = {10.1007/978-1-4684-5424-6_6},
   url = {https://dx.doi.org/10.1007/978-1-4684-5424-6_6},
   year = {1988}
}

@article{RN103,
   author = {Heindl, Jason E. and Crosby, Daniel and Brar, Sukhdev and Pinto, John F. and Singletary, Tiyan and Merenich, Daniel and Eagan, Justin L. and Buechlein, Aaron M. and Bruger, Eric L. and Waters, Christopher M. and Fuqua, Clay},
   title = {Reciprocal control of motility and biofilm formation by the PdhS2 two-component sensor kinase of \textit{Agrobacterium tumefaciens}},
   journal = {Microbiology},
   volume = {165},
   number = {2},
   pages = {146-162},
   ISSN = {1350-0872},
   DOI = {10.1099/mic.0.000758},
   url = {https://dx.doi.org/10.1099/mic.0.000758},
   year = {2019},
   type = {Journal Article}
}

@article{RN5,
   author = {Hornung, B. and Martins Dos Santos, V. A. P. and Smidt, H. and Schaap, P. J.},
   title = {Studying microbial functionality within the gut ecosystem by systems biology},
   journal = {Genes Nutr},
   volume = {13},
   pages = {5},
   ISSN = {1555-8932 (Print)
1555-8932 (Linking)},
   DOI = {10.1186/s12263-018-0594-6},
   url = {https://www.ncbi.nlm.nih.gov/pubmed/29556373},
   year = {2018}
}

@article{RN8,
   author = {Howard-Varona, Cristina and Hargreaves, Katherine R and Abedon, Stephen T and Sullivan, Matthew B},
   title = {Lysogeny in nature: mechanisms, impact and ecology of temperate phages},
   journal = {The ISME journal},
   volume = {11},
   number = {7},
   pages = {1511-1520},
   ISSN = {1751-7370},
   year = {2017}
}

@article{RN28,
   author = {Hu, J. and Ye, H. and Wang, S. and Wang, J. and Han, D.},
   title = {Prophage Activation in the Intestine: Insights Into Functions and Possible Applications},
   journal = {Front Microbiol},
   volume = {12},
   pages = {785634},
   ISSN = {1664-302X (Print)
1664-302X (Linking)},
   DOI = {10.3389/fmicb.2021.785634},
   url = {https://www.ncbi.nlm.nih.gov/pubmed/34966370},
   year = {2021}
}

@article{RN54,
   author = {Hurwitz, Bonnie L. and Deng, Li and Poulos, Bonnie T. and Sullivan, Matthew B.},
   title = {Evaluation of methods to concentrate and purify ocean virus communities through comparative, replicated metagenomics},
   journal = {Environmental Microbiology},
   volume = {15},
   number = {5},
   pages = {1428-1440},
   ISSN = {1462-2912},
   DOI = {10.1111/j.1462-2920.2012.02836.x},
   url = {https://dx.doi.org/10.1111/j.1462-2920.2012.02836.x},
   year = {2013},
   type = {Journal Article}
}

@article{RN76,
   author = {Isenberg, Ruth Y and Christensen, David G and Visick, Karen L and Mandel, Mark J},
   title = {High levels of cyclic diguanylate interfere with beneficial bacterial colonization},
   journal = {Mbio},
   volume = {13},
   number = {4},
   pages = {e01671-22},
   ISSN = {2150-7511},
   year = {2022},
   type = {Journal Article}
}

@article{RN75,
   author = {Jenal, Urs and Malone, Jacob},
   title = {Mechanisms of Cyclic-di-GMP Signaling in Bacteria},
   journal = {Annual Review of Genetics},
   volume = {40},
   number = {1},
   pages = {385-407},
   ISSN = {0066-4197},
   DOI = {10.1146/annurev.genet.40.110405.090423},
   year = {2006}
}

@article{RN13,
   author = {Jiang, S. C. and Paul, J. H.},
   title = {Significance of Lysogeny in the Marine Environment: Studies with Isolates and a Model of Lysogenic Phage Production},
   journal = {Microbial Ecology},
   volume = {35},
   number = {3},
   pages = {235-243},
   ISSN = {0095-3628},
   DOI = {10.1007/s002489900079},
   year = {1998}
}

@article{RN2,
   author = {Johansson, M. E. and Larsson, J. M. and Hansson, G. C.},
   title = {The two mucus layers of colon are organized by the MUC2 mucin, whereas the outer layer is a legislator of host-microbial interactions},
   journal = {Proc Natl Acad Sci U S A},
   volume = {108 Suppl 1},
   pages = {4659-65},
   ISSN = {1091-6490 (Electronic)
0027-8424 (Linking)},
   DOI = {10.1073/pnas.1006451107},
   url = {http://www.ncbi.nlm.nih.gov/pubmed/20615996},
   year = {2011}
}

@inbook{RN105,
   author = {Jones, Christopher J. and Wozniak, Daniel J.},
   title = {Congo Red Stain Identifies Matrix Overproduction and Is an Indirect Measurement for c-di-GMP in Many Species of Bacteria},
   booktitle = {c-di-GMP Signaling: Methods and Protocols},
   editor = {Sauer, Karin},
   publisher = {Springer New York},
   address = {New York, NY},
   pages = {147-156},
   ISBN = {978-1-4939-7240-1},
   DOI = {10.1007/978-1-4939-7240-1_12},
   url = {https://doi.org/10.1007/978-1-4939-7240-1_12},
   year = {2017},
   type = {Book Section}
}

@misc{RN69,
   author = {Kassambara, Alboukadel},
   title = {ggpubr: 'ggplot2' Based Publication Ready Plots},
   url = { 	https://rpkgs.datanovia.com/ggpubr/},
   year = {2020}
}

@article{RN58,
   author = {Kearns, D. B.},
   title = {A field guide to bacterial swarming motility},
   journal = {Nat Rev Microbiol},
   volume = {8},
   number = {9},
   pages = {634-44},
   ISSN = {1740-1534 (Electronic)
1740-1526 (Linking)},
   DOI = {10.1038/nrmicro2405},
   url = {https://www.ncbi.nlm.nih.gov/pubmed/20694026},
   year = {2010}
}

@article{RN48,
   author = {Kieft, Kristopher and Zhou, Zhichao and Anantharaman, Karthik},
   title = {VIBRANT: automated recovery, annotation and curation of microbial viruses, and evaluation of viral community function from genomic sequences},
   journal = {Microbiome},
   volume = {8},
   number = {1},
   ISSN = {2049-2618},
   DOI = {10.1186/s40168-020-00867-0},
   url = {https://dx.doi.org/10.1186/s40168-020-00867-0},
   year = {2020},
   type = {Journal Article}
}

@article{RN16,
   author = {Kim, Min-Soo and Bae, Jin-Woo},
   title = {Lysogeny is prevalent and widely distributed in the murine gut microbiota},
   journal = {The ISME Journal},
   volume = {12},
   number = {4},
   pages = {1127-1141},
   ISSN = {1751-7362},
   DOI = {10.1038/s41396-018-0061-9},
   url = {https://dx.doi.org/10.1038/s41396-018-0061-9},
   year = {2018}
}

@article{RN112,
   author = {Kovach, M. E. and Phillips, R. W. and Elzer, P. H. and Roop, R. M., 2nd and Peterson, K. M.},
   title = {pBBR1MCS: a broad-host-range cloning vector},
   journal = {Biotechniques},
   volume = {16},
   number = {5},
   pages = {800-2},
   keywords = {Base Sequence
Cloning, Molecular
*Genetic Vectors
Molecular Sequence Data
*Plasmids},
   ISSN = {0736-6205 (Print)
0736-6205},
   year = {1994},
   type = {Journal Article}
}


@phdthesis{RN114,
   author = {Leigh, B.},
   title = {Role of viruses within metaorganisms: \textit{Ciona intestinalis} as a model system},
   university = {University of South Florida},
   year = {2017},
   type = {Thesis}
}

@article{RN33,
   author = {Leigh, Brittany and Karrer, Charlotte and Cannon, John P and Breitbart, Mya and Dishaw, Larry J},
   title = {Isolation and characterization of a Shewanella phage–host system from the gut of the tunicate, \textit{Ciona intestinalis}},
   journal = {Viruses},
   volume = {9},
   number = {3},
   pages = {60},
   year = {2017}
}

@article{RN15,
   author = {Leigh, Brittany A. and Djurhuus, Anni and Breitbart, Mya and Dishaw, Larry J.},
   title = {The gut virome of the protochordate model organism, \textit{Ciona intestinalis} subtype A},
   journal = {Virus Research},
   volume = {244},
   pages = {137-146},
   keywords = {Gut
Microbiome
Viral metagenome
Phage},
   ISSN = {0168-1702},
   DOI = {https://doi.org/10.1016/j.virusres.2017.11.015},
   url = {http://www.sciencedirect.com/science/article/pii/S0168170217305117},
   year = {2018}
}

@article{RN60,
   author = {Leigh, B. A. and Liberti, A. and Dishaw, L. J.},
   title = {Generation of Germ-Free \textit{Ciona intestinalis} for Studies of Gut-Microbe Interactions},
   journal = {Front Microbiol},
   volume = {7},
   pages = {2092},
   keywords = {Ciona intestinalis
Gut colonization
germ-free
host-microbe interaction
microbiome},
   ISSN = {1664-302X (Print)
1664-302X (Linking)},
   DOI = {10.3389/fmicb.2016.02092},
   url = {https://www.ncbi.nlm.nih.gov/pubmed/28082961},
   year = {2016}
}

@article{RN1,
   author = {Li, H. and Limenitakis, J. P. and Fuhrer, T. and Geuking, M. B. and Lawson, M. A. and Wyss, M. and Brugiroux, S. and Keller, I. and Macpherson, J. A. and Rupp, S. and Stolp, B. and Stein, J. V. and Stecher, B. and Sauer, U. and McCoy, K. D. and Macpherson, A. J.},
   title = {The outer mucus layer hosts a distinct intestinal microbial niche},
   journal = {Nat Commun},
   volume = {6},
   pages = {8292},
   ISSN = {2041-1723 (Electronic)
2041-1723 (Linking)},
   DOI = {10.1038/ncomms9292},
   url = {http://www.ncbi.nlm.nih.gov/pubmed/26392213},
   year = {2015}
}

@article{RN38,
   author = {Liberti, A. and Cannon, J. P. and Litman, G. W. and Dishaw, L. J.},
   title = {A Soluble Immune Effector Binds Both Fungi and Bacteria via Separate Functional Domains},
   journal = {Front Immunol},
   volume = {10},
   pages = {369},
   ISSN = {1664-3224 (Electronic)
1664-3224 (Linking)},
   DOI = {10.3389/fimmu.2019.00369},
   url = {http://www.ncbi.nlm.nih.gov/pubmed/30894858},
   year = {2019}
}

@article{RN115,
   author = {Liberti, Assunta and Leigh, Brittany and De Santis, Rosaria and Pinto, Maria Rosaria and Cannon, John P and Dishaw, Larry J and Litman, Gary W},
   title = {An immune effector system in the protochordate gut sheds light on fundamental aspects of vertebrate immunity},
   journal = {Pathogen-Host Interactions: Antigenic Variation v. Somatic Adaptations},
   pages = {159-173},
   ISSN = {3319208187},
   year = {2015}
}

@inbook{RN56,
   author = {Liberti, Assunta and Leigh, Brittany A. and Graham, Zachary and Natarajan, Ojas and Dishaw, Larry J.},
   title = {A Role for Secreted Immune Effectors in Microbial Biofilm Formation Revealed by Simple In Vitro Assays},
   booktitle = {Immune Receptors: Methods and Protocols},
   editor = {Rast, Jonathan and Buckley, Katherine},
   publisher = {Springer US},
   address = {New York, NY},
   pages = {127-140},
   ISBN = {978-1-0716-1944-5},
   DOI = {10.1007/978-1-0716-1944-5_9},
   url = {https://doi.org/10.1007/978-1-0716-1944-5_9},
   year = {2022},
   type = {Book Section}
}

@article{RN37,
   author = {Liberti, Assunta and Natarajan, Ojas and Atkinson, Celine Grace F. and Sordino, Paolo and Dishaw, Larry J.},
   title = {Reflections on the Use of an Invertebrate Chordate Model System for Studies of Gut Microbial Immune Interactions},
   journal = {Frontiers in Immunology},
   volume = {12},
   ISSN = {1664-3224},
   DOI = {10.3389/fimmu.2021.642687},
   year = {2021}
}

@article{RN110,
   author = {Liberti, A. and Pollastro, C. and Pinto, G. and Illiano, A. and Marino, R. and Amoresano, A. and Spagnuolo, A. and Sordino, P.},
   title = {Transcriptional and proteomic analysis of the innate immune response to microbial stimuli in a model invertebrate chordate},
   journal = {Front Immunol},
   volume = {14},
   pages = {1217077},
   ISSN = {1664-3224},
   DOI = {10.3389/fimmu.2023.1217077},
   year = {2023}
}

@article{RN91,
   author = {Liberti, Assunta and Zucchetti, Ivana and Melillo, Daniela and Skapura, Diana and Shibata, Yoshimi and De Santis, Rosaria and Pinto, Maria Rosaria and Litman, Gary W. and Dishaw, Larry J.},
   title = {Chitin protects the gut epithelial barrier in a protochordate model of DSS-induced colitis},
   journal = {Biology Open},
   volume = {7},
   number = {1},
   ISSN = {2046-6390},
   DOI = {10.1242/bio.029355},
   url = {https://doi.org/10.1242/bio.029355},
   year = {2018}
}

@article{RN96,
   author = {Lin, Wei and Fullner, Karla Jean and Clayton, Rebecca and Sexton, Jessica A. and Rogers, Marc B. and Calia, Kerstin E. and Calderwood, Stephen B. and Fraser, Claire and Mekalanos, John J.},
   title = {Identification of a \textit{Vibrio cholerae} RTX toxin gene cluster that is tightly linked to the cholera toxin prophage},
   journal = {Proceedings of the National Academy of Sciences},
   volume = {96},
   number = {3},
   pages = {1071-1076},
   ISSN = {0027-8424},
   DOI = {10.1073/pnas.96.3.1071},
   url = {https://dx.doi.org/10.1073/pnas.96.3.1071},
   year = {1999}
}

@article{RN36,
   author = {Liu, Li-Ping and Xiang, Jian-Hai and Dong, Bo and Natarajan, Pavanasam and Yu, Kui-Jie and Cai, Nan-Er},
   title = {\textit{Ciona intestinalis} as an emerging model organism: Its regeneration under controlled conditions and methodology for egg dechorionation},
   journal = {Journal of Zhejiang University SCIENCE B},
   volume = {7},
   number = {6},
   pages = {467-474},
   ISSN = {1673-1581},
   DOI = {10.1631/jzus.2006.b0467},
   url = {https://dx.doi.org/10.1631/jzus.2006.b0467},
   year = {2006}
}

@article{RN30,
   author = {Liu, Xiaoxiao and Li, Yangmei and Guo, Yunxue and Zeng, Zhenshun and Li, Baiyuan and Wood, Thomas K. and Cai, Xingsheng and Wang, Xiaoxue},
   title = {Physiological Function of Rac Prophage During Biofilm Formation and Regulation of Rac Excision in \textit{Escherichia coli} K-12},
   journal = {Scientific Reports},
   volume = {5},
   number = {1},
   pages = {16074},
   ISSN = {2045-2322},
   DOI = {10.1038/srep16074},
   url = {https://dx.doi.org/10.1038/srep16074},
   year = {2015}
}

@article{RN43,
   author = {Liu, Xiaoxiao and Tang, Kaihao and Zhang, Dali and Li, Yangmei and Liu, Zhe and Yao, Jianyun and Wood, Thomas K. and Wang, Xiaoxue},
   title = {Symbiosis of a P2‐family phage and deep‐sea \textit{Shewanella putrefaciens}},
   journal = {Environmental Microbiology},
   volume = {21},
   number = {11},
   pages = {4212-4232},
 
   ISSN = {1462-2912},
   DOI = {10.1111/1462-2920.14781},
   url = {https://dx.doi.org/10.1111/1462-2920.14781},
   year = {2019}
}

@article{RN68,
   author = {Lord, Samuel J and Velle, Katrina B and Mullins, R Dyche and Fritz-Laylin, Lillian K},
   title = {SuperPlots: Communicating reproducibility and variability in cell biology},
   journal = {Journal of Cell Biology},
   volume = {219},
   number = {6},
   ISSN = {0021-9525},
   year = {2020}
}

@article{RN9,
   author = {Lwoff, André},
   title = {Lysogeny},
   journal = {Bacteriological reviews},
   volume = {17},
   number = {4},
   pages = {269-337},
   ISSN = {0005-3678},
   year = {1953}
}

@article{RN79,
   author = {Lynch, Jonathan B. and Bennett, Brittany D. and Merrill, Bryan D. and Ruby, Edward G. and Hryckowian, Andrew J.},
   title = {Independent host- and bacterium-based determinants protect a model symbiosis from phage predation},
   journal = {Cell Reports},
   volume = {38},
   number = {7},
   pages = {110376},
   keywords = {bacteriophages
symbiosis
microbe-host interactions
bacterial genetics
genomic analysis
model systems},
   ISSN = {2211-1247},
   DOI = {https://doi.org/10.1016/j.celrep.2022.110376},
   url = {https://www.sciencedirect.com/science/article/pii/S2211124722000973},
   year = {2022},
   type = {Journal Article}
}

@article{RN23,
   author = {Maiques, E. and Ubeda, C. and Campoy, S. and Salvador, N. and Lasa, I. and Novick, R. P. and Barbe, J. and Penades, J. R.},
   title = {beta-lactam antibiotics induce the SOS response and horizontal transfer of virulence factors in Staphylococcus aureus},
   journal = {J Bacteriol},
   volume = {188},
   number = {7},
   pages = {2726-9},
   ISSN = {0021-9193 (Print)
0021-9193 (Linking)},
   DOI = {10.1128/JB.188.7.2726-2729.2006},
   url = {http://www.ncbi.nlm.nih.gov/pubmed/16547063},
   year = {2006}
}

@article{RN72,
   author = {Männistö, Riina H. and Kivelä, Hanna M. and Paulin, Lars and Bamford, Dennis H. and Bamford, Jaana K.H.},
   title = {The Complete Genome Sequence of PM2, the First Lipid-Containing Bacterial Virus To Be Isolated},
   journal = {Virology},
   volume = {262},
   number = {2},
   pages = {355-363},
   ISSN = {0042-6822},
   DOI = {10.1006/viro.1999.9837},
   year = {1999},
   type = {Journal Article}
}

@article{RN116,
   author = {Marino, Rita and Melillo, Daniela and Italiani, Paola and Boraschi, Diana},
   title = {Environmental stress and nanoplastics’ effects on \textit{Ciona robusta}: regulation of immune/stress-related genes and induction of innate memory in pharynx and gut},
   journal = {Frontiers in Immunology},
   volume = {14},
   pages = {1176982},
   ISSN = {1664-3224},
   year = {2023}
}

@article{RN62,
   author = {Miles, Ashley A and Misra, SS and Irwin, JO},
   title = {The estimation of the bactericidal power of the blood},
   journal = {Epidemiology \& Infection},
   volume = {38},
   number = {6},
   pages = {732-749},
   ISSN = {1469-4409},
   year = {1938}
}

@article{RN11,
   author = {Mills, S. and Shanahan, F. and Stanton, C. and Hill, C. and Coffey, A. and Ross, R. P.},
   title = {Movers and shakers: influence of bacteriophages in shaping the mammalian gut microbiota},
   journal = {Gut Microbes},
   volume = {4},
   number = {1},
   pages = {4-16},
   ISSN = {1949-0984 (Electronic)},
   DOI = {10.4161/gmic.22371},
   url = {http://www.ncbi.nlm.nih.gov/pubmed/23022738},
   year = {2013}
}

@article{RN86,
   author = {Minot, S. and Bryson, A. and Chehoud, C. and Wu, G. D. and Lewis, J. D. and Bushman, F. D.},
   title = {Rapid evolution of the human gut virome},
   journal = {Proc Natl Acad Sci U S A},
   volume = {110},
   number = {30},
   pages = {12450-5},
   ISSN = {1091-6490 (Electronic)
0027-8424 (Linking)},
   DOI = {10.1073/pnas.1300833110},
   url = {http://www.ncbi.nlm.nih.gov/pubmed/23836644},
   year = {2013}
}

@article{RN85,
   author = {Minot, S. and Sinha, R. and Chen, J. and Li, H. and Keilbaugh, S. A. and Wu, G. D. and Lewis, J. D. and Bushman, F. D.},
   title = {The human gut virome: inter-individual variation and dynamic response to diet},
   journal = {Genome Res},
   volume = {21},
   number = {10},
   pages = {1616-25},
   ISSN = {1549-5469 (Electronic)
1088-9051 (Linking)},
   DOI = {10.1101/gr.122705.111},
   url = {http://www.ncbi.nlm.nih.gov/pubmed/21880779},
   year = {2011}
}

@article{RN7,
   author = {Mirzaei, Mohammadali Khan and Maurice, Corinne F.},
   title = {Ménage à trois in the human gut: interactions between host, bacteria and phages},
   journal = {Nature Reviews Microbiology},
   volume = {15},
   number = {7},
   pages = {397-408},
   ISSN = {1740-1526},
   DOI = {10.1038/nrmicro.2017.30},
   url = {https://dx.doi.org/10.1038/nrmicro.2017.30},
   year = {2017}
}

@article{RN6,
   author = {Moran, N. A. and Ochman, H. and Hammer, T. J.},
   title = {Evolutionary and ecological consequences of gut microbial communities},
   journal = {Annu Rev Ecol Evol Syst},
   volume = {50},
   number = {1},
   pages = {451-475},
   ISSN = {1543-592X (Print)
1543-592X (Linking)},
   DOI = {10.1146/annurev-ecolsys-110617-062453},
   url = {https://www.ncbi.nlm.nih.gov/pubmed/32733173},
   year = {2019}
}

@article{RN24,
   author = {Nanda, A. M. and Thormann, K. and Frunzke, J.},
   title = {Impact of spontaneous prophage induction on the fitness of bacterial populations and host-microbe interactions},
   journal = {J Bacteriol},
   volume = {197},
   number = {3},
   pages = {410-9},
   ISSN = {1098-5530 (Electronic)
0021-9193 (Linking)},
   DOI = {10.1128/JB.02230-14},
   url = {http://www.ncbi.nlm.nih.gov/pubmed/25404701},
   year = {2015}
}

@misc{RN47,
   author = {Parks, Donovan},
   title = {CompareM: Comparative Genomic Statistics},
   url = {https://github.com/dparks1134/CompareM},
   year = {2020},
   type = {Computer Program}
}

@article{RN46,
   author = {Parks, D. H. and Imelfort, M. and Skennerton, C. T. and Hugenholtz, P. and Tyson, G. W.},
   title = {CheckM: assessing the quality of microbial genomes recovered from isolates, single cells, and metagenomes},
   journal = {Genome Res},
   volume = {25},
   number = {7},
   pages = {1043-55},
   keywords = {*Genome, Microbial
*Metagenome
Metagenomics/*methods},
   ISSN = {1549-5469 (Electronic)
1088-9051 (Linking)},
   DOI = {10.1101/gr.186072.114},
   url = {https://www.ncbi.nlm.nih.gov/pubmed/25977477},
   year = {2015}
}

@article{RN64,
   author = {Pfaffl, M. W.},
   title = {A new mathematical model for relative quantification in real-time RT-PCR},
   journal = {Nucleic Acids Res},
   volume = {29},
   number = {9},
   pages = {e45},
   ISSN = {1362-4962 (Electronic)
0305-1048 (Linking)},
   DOI = {10.1093/nar/29.9.e45},
   url = {https://www.ncbi.nlm.nih.gov/pubmed/11328886},
   year = {2001}
}

@article{RN98,
   author = {Pfaffl, Michael W and Tichopad, Ales and Prgomet, Christian and Neuvians, Tanja P},
   title = {Determination of stable housekeeping genes, differentially regulated target genes and sample integrity: BestKeeper–Excel-based tool using pair-wise correlations},
   journal = {Biotechnology letters},
   volume = {26},
   pages = {509-515},
   ISSN = {0141-5492},
   year = {2004}
}

@article{RN82,
   author = {Pratt, L. A. and Kolter, R.},
   title = {Genetic analysis of \textit{Escherichia coli} biofilm formation: roles of flagella, motility, chemotaxis and type I pili},
   journal = {Mol Microbiol},
   volume = {30},
   number = {2},
   pages = {285-93},
   ISSN = {0950-382X (Print)
0950-382X (Linking)},
   url = {http://www.ncbi.nlm.nih.gov/pubmed/9791174},
   year = {1998}
}

@article{RN87,
   author = {Reyes, A. and Haynes, M. and Hanson, N. and Angly, F. E. and Heath, A. C. and Rohwer, F. and Gordon, J. I.},
   title = {Viruses in the faecal microbiota of monozygotic twins and their mothers},
   journal = {Nature},
   volume = {466},
   number = {7304},
   pages = {334-8},
   ISSN = {1476-4687 (Electronic)
0028-0836 (Linking)},
   DOI = {10.1038/nature09199},
   url = {http://www.ncbi.nlm.nih.gov/pubmed/20631792},
   year = {2010}
}

@article{RN31,
   author = {Rice, Scott A. and Tan, Chuan Hao and Mikkelsen, Per Jensen and Kung, Vanderlene and Woo, Jerry and Tay, Martin and Hauser, Alan and McDougald, Diane and Webb, Jeremy S. and Kjelleberg, Staffan},
   title = {The biofilm life cycle and virulence of \textit{Pseudomonas aeruginosa} are dependent on a filamentous prophage},
   journal = {The ISME Journal},
   volume = {3},
   number = {3},
   pages = {271-282},
   ISSN = {1751-7362},
   DOI = {10.1038/ismej.2008.109},
   url = {https://dx.doi.org/10.1038/ismej.2008.109},
   year = {2009}
}

@article{RN107,
   author = {Roberts, Steven and Gueguen, Yannick and de Lorgeril, Julien and Goetz, Frederick},
   title = {Rapid accumulation of an interleukin 17 homolog transcript in \textit{Crassostrea gigas} hemocytes following bacterial exposure},
   journal = {Developmental \& Comparative Immunology},
   volume = {32},
   number = {9},
   pages = {1099-1104},
   ISSN = {0145-305X},
   DOI = {https://doi.org/10.1016/j.dci.2008.02.006},
   url = {https://www.sciencedirect.com/science/article/pii/S0145305X08000451},
   year = {2008}
}

@article{RN74,
   author = {Rooks, Michelle G. and Garrett, Wendy S.},
   title = {Gut microbiota, metabolites and host immunity},
   journal = {Nature Reviews Immunology},
   volume = {16},
   number = {6},
   pages = {341-352},
   ISSN = {1474-1733},
   DOI = {10.1038/nri.2016.42},
   url = {https://dx.doi.org/10.1038/nri.2016.42},
   year = {2016}
}

@article{RN49,
   author = {Roux, Simon and Enault, Francois and Hurwitz, Bonnie L. and Sullivan, Matthew B.},
   title = {VirSorter: mining viral signal from microbial genomic data},
   journal = {PeerJ},
   volume = {3},
   pages = {e985},
   ISSN = {2167-8359},
   DOI = {10.7717/peerj.985},
   url = {https://dx.doi.org/10.7717/peerj.985},
   year = {2015},
   type = {Journal Article}
}

@article{RN51,
   author = {Saltikov, Chad W. and Newman, Dianne K.},
   title = {Genetic identification of a respiratory arsenate reductase},
   journal = {Proceedings of the National Academy of Sciences},
   volume = {100},
   number = {19},
   pages = {10983-10988},
   DOI = {10.1073/pnas.1834303100},
   url = {https://www.pnas.org/content/pnas/100/19/10983.full.pdf},
   year = {2003}
}

@article{RN17,
   author = {Shkoporov, Andrey N. and Hill, Colin},
   title = {Bacteriophages of the Human Gut: The “Known Unknown” of the Microbiome},
   journal = {Cell Host \& Microbe},
   volume = {25},
   number = {2},
   pages = {195-209},
   ISSN = {1931-3128},
   DOI = {10.1016/j.chom.2019.01.017},
   url = {https://dx.doi.org/10.1016/j.chom.2019.01.017},
   year = {2019}
}

@article{RN14,
   author = {Silveira, Cynthia B. and Luque, Antoni and Rohwer, Forest},
   title = {The landscape of lysogeny across microbial community density, diversity and energetics},
   journal = {Environmental Microbiology},
   volume = {23},
   number = {8},
   pages = {4098-4111},
   keywords = {Literature review
lysogeny},
   ISSN = {1462-2912},
   DOI = {10.1111/1462-2920.15640},
   url = {https://dx.doi.org/10.1111/1462-2920.15640},
   year = {2021}
}

@article{RN29,
   author = {Swidsinski, A. and Loening-Baucke, V. and Theissig, F. and Engelhardt, H. and Bengmark, S. and Koch, S. and Lochs, H. and Dorffel, Y.},
   title = {Comparative study of the intestinal mucus barrier in normal and inflamed colon},
   journal = {Gut},
   volume = {56},
   number = {3},
   pages = {343-50},
   ISSN = {0017-5749 (Print)
0017-5749 (Linking)},
   DOI = {10.1136/gut.2006.098160},
   url = {http://www.ncbi.nlm.nih.gov/pubmed/16908512},
   year = {2007}
}

@article{RN77,
   author = {Tan, Demeng and Hansen, Mads Frederik and De Carvalho, Luís Nunes and Røder, Henriette Lyng and Burmølle, Mette and Middelboe, Mathias and Svenningsen, Sine Lo},
   title = {High cell densities favor lysogeny: induction of an H20 prophage is repressed by quorum sensing and enhances biofilm formation in \textit{Vibrio anguillarum}},
   journal = {The ISME Journal},
   volume = {14},
   number = {7},
   pages = {1731-1742},
   ISSN = {1751-7362},
   DOI = {10.1038/s41396-020-0641-3},
   url = {https://dx.doi.org/10.1038/s41396-020-0641-3},
   year = {2020},
   type = {Journal Article}
}

@inbook{RN117,
   author = {Taylor, Véronique L. and Fitzpatrick, Alexa D. and Islam, Zafrin and Maxwell, Karen L.},
   title = {Chapter One - The Diverse Impacts of Phage Morons on Bacterial Fitness and Virulence},
   booktitle = {Advances in Virus Research},
   editor = {Kielian, Margaret and Mettenleiter, Thomas C. and Roossinck, Marilyn J.},
   publisher = {Academic Press},
   volume = {103},
   pages = {1-31},
   ISBN = {0065-3527},
   DOI = {https://doi.org/10.1016/bs.aivir.2018.08.001},
   url = {https://www.sciencedirect.com/science/article/pii/S0065352718300435},
   year = {2019},
   type = {Book Section}
}

@misc{RN65,
   author = {Team, R Core},
   title = {R : A language and environment for statistical  computing.},
   publisher= {R Team},
   url = { https://www.R-project.org/},
   year = {2021}
}

@article{RN81,
   author = {Thomas, A. D. and Parker, W.},
   title = {Cultivation of epithelial-associated microbiota by the immune system},
   journal = {Future Microbiol},
   volume = {5},
   number = {10},
   pages = {1483-92},
   ISSN = {1746-0921 (Electronic)
1746-0913 (Linking)},
   DOI = {10.2217/fmb.10.108},
   url = {http://www.ncbi.nlm.nih.gov/pubmed/21073309},
   year = {2010}
}

@article{RN78,
   author = {Tsao, Yu-Fan and Taylor, Véronique L. and Kala, Smriti and Bondy-Denomy, Joseph and Khan, Alima N. and Bona, Diane and Cattoir, Vincent and Lory, Stephen and Davidson, Alan R. and Maxwell, Karen L.},
   title = {Phage Morons Play an Important Role in \textit{Pseudomonas aeruginosa} Phenotypes},
   journal = {Journal of Bacteriology},
   volume = {200},
   number = {22},
   pages = {e00189-18},
   DOI = {doi:10.1128/JB.00189-18},
   url = {https://journals.asm.org/doi/abs/10.1128/JB.00189-18},
   year = {2018}
}

@article{RN100,
   author = {Vandesompele, Jo and De Preter, Katleen and Pattyn, Filip and Poppe, Bruce and Van Roy, Nadine and De Paepe, Anne and Speleman, Frank},
   title = {Accurate normalization of real-time quantitative RT-PCR data by geometric averaging of multiple internal control genes},
   journal = {Genome biology},
   volume = {3},
   number = {7},
   pages = {1-12},
   ISSN = {1474-760X},
   year = {2002}
}

@article{RN90,
   author = {Wagner, P. L. and Waldor, M. K.},
   title = {Bacteriophage control of bacterial virulence},
   journal = {Infect Immun},
   volume = {70},
   number = {8},
   pages = {3985-93},
   ISSN = {0019-9567 (Print)
0019-9567 (Linking)},
   url = {http://www.ncbi.nlm.nih.gov/pubmed/12117903},
   year = {2002}
}

@article{RN25,
   author = {Wang, X. and Kim, Y. and Ma, Q. and Hong, S. H. and Pokusaeva, K. and Sturino, J. M. and Wood, T. K.},
   title = {Cryptic prophages help bacteria cope with adverse environments},
   journal = {Nat Commun},
   volume = {1},
   pages = {147},
   ISSN = {2041-1723 (Electronic)
2041-1723 (Linking)},
   DOI = {10.1038/ncomms1146},
   url = {http://www.ncbi.nlm.nih.gov/pubmed/21266997},
   year = {2010}
}

@article{RN92,
   author = {Watnick, Paula and Kolter, Roberto},
   title = {Biofilm, city of microbes},
   journal = {Journal of bacteriology},
   volume = {182},
   number = {10},
   pages = {2675-2679},
   ISSN = {1098-5530},
   year = {2000}
}

@article{RN88,
   author = {Weinbauer, M. G. and Suttle, C. A.},
   title = {Lysogeny and prophage induction in coastal and offshore bacterial communities},
   journal = {Aquatic Microbial Ecology},
   volume = {18},
   number = {3},
   pages = {217-225},
   ISSN = {0948-3055},
   DOI = {10.3354/ame018217},
   url = {<Go to ISI>://WOS:000082124500002},
   year = {1999}
}

@article{RN45,
   author = {Wick, Ryan R and Judd, Louise M and Gorrie, Claire L and Holt, Kathryn E},
   title = {Unicycler: resolving bacterial genome assemblies from short and long sequencing reads},
   journal = {PLoS computational biology},
   volume = {13},
   number = {6},
   pages = {e1005595},
   ISSN = {1553-734X},
   year = {2017}
}

@misc{RN66,
   author = {Wickham, Hadley},
   title = {ggplot2: Elegant Graphics for Data Analysis},
   ISBN = {978-3-319-24277-4},
   url = {https://ggplot2.tidyverse.org},
   year = {2016},
   type = {Computer Program}
}

@article{RN57,
   author = {Wolfe, A. J. and Berg, H. C.},
   title = {Migration of bacteria in semisolid agar},
   journal = {Proc Natl Acad Sci U S A},
   volume = {86},
   number = {18},
   pages = {6973-7},
   ISSN = {0027-8424 (Print)
0027-8424 (Linking)},
   DOI = {10.1073/pnas.86.18.6973},
   url = {https://www.ncbi.nlm.nih.gov/pubmed/2674941},
   year = {1989}
}

@article{RN97,
   author = {Xie, F. and Xiao, P. and Chen, D. and Xu, L. and Zhang, B.},
   title = {miRDeepFinder: a miRNA analysis tool for deep sequencing of plant small RNAs},
   journal = {Plant Mol Biol},
   ISSN = {0167-4412},
   DOI = {10.1007/s11103-012-9885-2},
   year = {2012}
}

@article{RN26,
   author = {Zhang, X. and McDaniel, A. D. and Wolf, L. E. and Keusch, G. T. and Waldor, M. K. and Acheson, D. W.},
   title = {Quinolone antibiotics induce Shiga toxin-encoding bacteriophages, toxin production, and death in mice},
   journal = {J Infect Dis},
   volume = {181},
   number = {2},
   pages = {664-70},
   ISSN = {0022-1899 (Print)
0022-1899 (Linking)},
   DOI = {10.1086/315239},
   url = {http://www.ncbi.nlm.nih.gov/pubmed/10669353},
   year = {2000}
}

@article{RN120,
   author = {Fraune, Sebastian and Bosch, Thomas CG},
   title = {Why bacteria matter in animal development and evolution},
   journal = {Bioessays},
   volume = {32},
   number = {7},
   pages = {571-580},
   ISSN = {0265-9247},
   year = {2010}
}
@ARTICLE{RN121,
  title     = "Bacteriophage {PM2} has a protein capsid surrounding a spherical
               proteinaceous lipid core",
  author    = "Kivel{\"a}, Hanna M and Kalkkinen, Nisse and Bamford, Dennis H",
  journal   = "J. Virol.",
  publisher = "American Society for Microbiology",
  volume    =  76,
  number    =  16,
  pages     = "8169--8178",
  year      =  2002,
  language  = "en"
}
