## Supplementary material for "Prophage regulation of *Shewanella fidelis* 3313 motility and biofilm formation: implications for gut colonization dynamics in *Ciona robusta*": Tex: elife-template.tex

%%%%%%%%%%%%%%%%%%%%%%%%%%%%%%%%%%%%%%%%%%%%%%%%%%%%%%%%%%%%
%%% ELIFE ARTICLE TEMPLATE
%%%%%%%%%%%%%%%%%%%%%%%%%%%%%%%%%%%%%%%%%%%%%%%%%%%%%%%%%%%%
%%% PREAMBLE 
\documentclass[10pt,lineno]{elife}
% Use the onehalfspacing option for 1.5 line spacing
% Use the doublespacing option for 2.0 line spacing
% Please note that these options may affect formatting.
% Additionally, the use of the \newcommand function should be limited.

\usepackage{newfile}

\usepackage{lipsum} % Required to insert dummy text
\usepackage[version=4]{mhchem}
\usepackage{siunitx}
\usepackage{caption} % Optional: for better caption control
\usepackage{booktabs} % For \toprule, \midrule, \bottomrule
\usepackage{float} % For [H] specifier
\usepackage{placeins} % For \FloatBarrier
\usepackage{tabularx}
\usepackage{longtable}
\usepackage{ragged2e}
\usepackage{booktabs}
\DeclareSIUnit\Molar{M}

%%%%%%%%%%%%%%%%%%%%%%%%%%%%%%%%%%%%%%%%%%%%%%%%%%%%%%%%%%%%
%%% ARTICLE SETUP
%%%%%%%%%%%%%%%%%%%%%%%%%%%%%%%%%%%%%%%%%%%%%%%%%%%%%%%%%%%%
\title{Prophage regulation of \textit{Shewanella fidelis} 3313 motility and biofilm formation: implications for gut colonization dynamics in \textit{\textit{Ciona} robusta}}

\author[1\authfn{2}]{Ojas Natarajan}
\author[1]{Susanne L. Gibboney}
\author[1]{Morgan N. Young}
\author[2]{Shen Jean Lim}
\author[3]{Felicia Nguyen}
\author[3\authfn{1}]{Natalia Pluta}
\author[4\authfn{4}]{Celine G.F. Atkinson}
\author[1,5]{Assunta Liberti}
\author[6\authfn{6}]{Eric D. Kees}
\author[1,2\authfn{7}]{Brittany A. Leigh}
\author[2]{Mya Breitbart}
\author[6,7]{Jeffrey A. Gralnick}
\author[1,3,4*]{Larry J. Dishaw}

\affil[1]{Department of Pediatrics, Morsani College of Medicine, University of South Florida, Tampa FL}
\affil[2]{College of Marine Science, University of South Florida, Tampa FL}
\affil[3]{Biomedical Sciences Program, University of South Florida, Tampa FL}
\affil[4]{Department of Cell Biology, Microbiology, and Molecular Biology}
\affil[5]{Stazione Zoologica Anton Dohrn, Naples, Italy}
\affil[6]{BioTechnology Institute, University of Minnesota, St. Paul, MN}
\affil[7]{Plant and Microbial Biology, University of Minnesota, St. Paul, MN USA}
\corr{}{LJD}

\presentadd[\authfn{1}]{Uniformed Services University of Health Sciences, Bethesda, MD, USA}
\presentadd[\authfn{4}]{School of Dentistry, University of Washington, Seattle, USA}
\presentadd[\authfn{6}]{Cargill Corporation, Wayzata, MN}
\presentadd[\authfn{7}]{LifeSci Communications, New York and Boston}


%%%%%%%%%%%%%%%%%%%%%%%%%%%%%%%%%%%%%%%%%%%%%%%%%%%%%%%%%%%%
%%% ARTICLE START
%%%%%%%%%%%%%%%%%%%%%%%%%%%%%%%%%%%%%%%%%%%%%%%%%%%%%%%%%%%%

\begin{document}


\maketitle


\begin{abstract}
Lysogens, bacteria with one or more viruses (prophages) integrated into their genomes, are abundant in the gut of animals. Prophages often influence bacterial traits; however, the influence of prophages on the gut microbiota-host immune axis in animals remains poorly understood. Here, we investigate the influence of the prophage SfPat on \textit{Shewanella fidelis} 3313, a persistent member of the gut microbiome of the model marine tunicate, \textit{\textit{Ciona} robusta}. Establishment of a SfPat deletion mutant ($\Delta$SfPat) reveals the influence of this prophage on bacterial physiology \textit{in vitro} and during colonization of the \textit{Ciona} gut. \textit{In vitro}, deletion of SfPat reduces \textit{S. fidelis} 3313 motility and swimming while increasing biofilm formation. To understand the \textit{in vivo} impact of these prophage-induced changes in bacterial traits, we exposed metamorphic stage 4 \textit{Ciona} juveniles to wildtype (WT) and $\Delta$SfPat strains. During colonization, $\Delta$SfPat localizes to overlapping and distinct areas of the gut compared to the WT strain. We examined the differential expression of various regulators of cyclic-di-GMP, a secondary signaling molecule that mediates biofilm formation and motility.  The \textit{pdeB} gene, which encodes a bacterial phosphodiesterase known to influence biofilm formation and motility by degrading cyclic-di-GMP, is upregulated in the WT strain but not in $\Delta$SfPat when examined \textit{in vivo}. Expression of the \textit{Ciona} gut immune effector, VCBP-C, is enhanced during colonization by $\Delta$SfPat compared to the WT strain; however, VCBP-C binding to the WT strain does not promote the excision of SfPat in an SOS-dependent pathway. Instead, VCBP-C binding significantly reduces the expression of a phage major capsid protein. Our findings suggest that SfPat influences host perception of this important colonizing commensal and highlights the significance of investigating tripartite dynamics between prophages, bacteria, and their animal hosts to better understand the gut microbiota-host immune axis.elife
\end{abstract}


\section{Introduction}

An epithelial layer with a mucus-rich surface lines the gastrointestinal tract (or gut) of animals. This dynamic environment is a primary interface between host immunity, symbiotic microbes, and dietary antigens \cite{RN1, RN2}. During colonization of the gut, bacteria encounter physical, chemical, and biological forces. They must compete for nutrients and niche space while managing the stress of digestive enzymes and host immune factors \cite{RN3, RN4, RN5, RN6}. Gut-colonizing bacteria also encounter bacteriophages (phages), or viruses that infect bacteria \cite{RN7}; e.g., over $10^{12}$ viruses have been estimated in the human gut \cite{RN17}.\\

Phages display both lytic and temperate lifestyles. While the former impacts bacterial community dynamics through lysis, the effect of temperate phages on bacterial communities, especially those associated with animals, remains poorly understood. Conventionally, temperate phages integrate into bacterial genomes as prophages and remain ‘dormant’ until an external trigger activates them to enter the lytic cycle \cite{RN9, RN10, RN8}; these bacteria are considered ‘lysogenized’ and referred to as lysogens \cite{RN9, RN8}. Prophages often encode accessory genes that can influence bacterial traits and behaviors \cite{RN11}. These genes can encode virulence factors, antibiotic resistance determinants, and those that provide superinfection exclusion, thereby protecting their bacterial hosts from infections by related phages \cite{RN12}. Based on the site of integration, prophages can also impact the expression of bacterial genes  \cite{RN95}. Bacterial lysogens exist in every environment \cite{RN13, RN14, RN15} and are particularly prevalent in the microbiomes of diverse animals \cite{RN16, RN17}.\\

Prophage induction results in bacterial lysis and can be mediated by various stressors, including antibiotics and inflammatory processes \cite{RN18,RN19, RN20,RN21,RN22,RN23, RN24, RN25, RN26}. Because prophages can influence the phenotypes of their bacterial hosts, such as biofilm formation \cite{RN24},  understanding the impact of lysogens in animal microbiomes is becoming a research priority \cite{RN27, RN28, RN96}. In the gut, biofilms that associate with host mucus may benefit the host by enhancing epithelial barriers against pathogenic invasion \cite{RN29}. Integration of prophages into bacterial genomes may impart functional changes that could be important in surface colonization. For example, in \textit{E. coli} K-12, integrating the Rac prophage into a tRNA thioltransferase region disrupts biofilm functions \cite{RN30}; deleting this prophage decreases resistance to antibiotics, acids and oxidative stress. Some of these traits may be affected by prophage-specific genes \cite{RN25}. Prophages have also been shown to influence biofilm life cycles in \textit{Pseudomonas aeruginosa} \cite{RN31}.\\

Lysogenized \textit{Shewanella} species colonize the gut of \textit{Ciona robusta}, an ascidian collected in Southern California waters \cite{RN32,RN33} and referred to here as {\textit{Ciona}}. Tunicates like \textit{Ciona} are a subphylum (Tunicata) of chordates that are well-established invertebrate model systems for studies of animal development \cite{RN34, RN35, RN36} and are now increasingly leveraged for gut immune and microbiome studies \cite{RN37}. Armed with only innate immunity, \textit{Ciona} maintains stable gut bacterial and viral communities \cite{RN32, RN15} despite continuously filtering microbe-rich seawater. Previous efforts to define gut immunity in \textit{Ciona} revealed the presence of a secreted immune effector, the variable immunoglobulin (V-Ig) domain-containing chitin-binding protein, or VCBP, that likely plays important roles in shaping the ecology of the gut microbiome by binding bacteria and fungi (as well as chitin-rich mucus) on opposing functional domains \cite{RN41, RN38}. VCBP-C is one of the best-studied VCBPs expressed in the stomach and intestines of \textit{Ciona}, shown to bind bacteria \textit{in vitro} and in the gut lumen \cite{RN40,RN41}. Based on various \textit{in vitro} and \textit{in vivo} observations, it was proposed previously that VCBPs likely modulate bacterial settlement and/or biofilm formation \cite{RN40,RN38,RN37}. The potential influence of soluble immune effectors on host-bacterial-viral interactions is particularly interesting. However, the possibility that prophages may influence interactions between bacteria and secreted immune effectors like VCBPs remains to be explored. \\

\textit{Shewanella fidelis} strain 3313 was isolated previously from the gut of \textit{Ciona} and found to posses two inducible prophages, SfPat and SfMu \cite{RN33}. Furthermore, \textit{in vitro} experiments demonstrated enhanced biofilm formation in \textit{S. fidelis} 3313 in the presence of extracellular DNA (eDNA) that may originate from lytic phage activity \cite{RN33}. Other \textit{Shewanella} species have previously demonstrated a link between phage-mediated lysis and biofilm formation \cite{RN42}. For example, in \textit{S. oneidensis} strain MR-1, Mu, and Lambda prophages enhance biofilm formation via eDNA released during prophage-induced lysis, with genomic DNA likely serving as a scaffold for biofilms \cite{RN42}. Similarly, the P2 prophage of \textit{S. putrefaciens} strain W3-18-1 influences biofilm formation via spontaneous induction at low frequencies, resulting in cell lysis and contributing eDNA that can mediate biofilm formation \cite{RN43}.\\

Here, we set out to isolate and characterize the influence of the prophage, SfPat, on its host, \textit{S. fidelis} 3313. Since the last description \cite{RN33}, the genome of \textit{S. fidelis} 3313 was improved by combining long-read and short-read sequencing, which resulted in improved resolution of the genomic landscape within and around the prophages. A homologous recombination-based deletion strategy was designed to generate a deletion mutant (i.e., knockout) of SfPat. We report that deletion of SfPat results in reduced bacterial motility and increased biofilm formation \textit{in vitro}. These changes in bacterial traits and behaviors are associated with the expression of genes regulating important signaling molecules and a corresponding impact on host immune gene expression during gut colonization in \textit{Ciona} juveniles. Gut colonization experiments in laboratory-reared \textit{Ciona} juveniles comparing wild-type (WT) and SfPat prophage knockout ($\Delta$SfPat) mutant strains demonstrate that SfPat influences gut colonization outcomes, e.g., niche preference and retention. These effects are influenced by host gene expression. The results reported herein reflect complex tripartite interactions among prophages, bacterial hosts, and animal immune systems. \\

\section{Results}

\subsection{Sequence verification of prophage deletion mutant strains}

Colony PCR and single primer extension sequencing were both used to validate the prophage deletion, using primers EDK81/82 for SfPat (\nameref{tab:deletelist})(\nameref{fig:1} a). All recovered amplicon sizes were consistent with the predictions for SfPat deletion (\nameref{fig:1} b). The SfPat deletion ($\Delta$SfPat) strain was named JG3862 (\nameref{tab:strainslist}). Genome sequencing of the deletion mutant strain did not reveal the significant introduction of additional mutations or DNA modifications (\nameref{fig:1} b).  The WT and $\Delta$SfPat strains were then used for \textit{in vitro} and \textit{in vivo} experiments to understand the potential role of prophages in shaping \textit{S. fidelis} 3313 colonization dynamics in the gut of \textit{Ciona}. \\
\begin{figure}[h]
\begin{fullwidth}
  \includegraphics[width=\textwidth]{Figures/Figure 1.pdf}
  \caption[Figure 1] {General prophage deletion scheme. (a) Location of upstream, downstream, and flanking primers used in the deletion of SfPat, primer orientation shown with respect to the prophage. (b) Deletion of SfPat from \textit{S. fidelis} 3313 identified after assembling Illumina (short-read) sequenced genomes and mapping onto the improved (short and long-read, PacBio, sequencing) WT genome. The figure illustrates SfPat deletion as revealed by subsequent Illumina sequencing. The solid gray areas on the SfPat genome indicate regions that share identity with the WT.}
  \label{fig:1}
\end{fullwidth} 
\end{figure}

\subsection{Prophage deletion modulates biofilm formation and motility in \textit{S. fidelis} 3313 \textit{in vitro}}

Deletion of SfPat from \textit{S. fidelis} 3313 contributed to an overall increase in biofilm formation as quantified by crystal violet staining by over 14\% compared to WT strains (\nameref{fig:2} a). Conventionally, to form a biofilm, bacteria will settle and initiate stationary growth dynamics \cite{RN92}. We studied bacterial swimming on simple semi-solid media to determine if the prophage influenced swimming motility in \textit{S. fidelis} 3313.  Bacterial motility was measured by the spread diameter from a primary inoculation point after overnight incubation (\nameref{fig:2} b). The WT strain demonstrated a mean diameter of 8.21 mm, while $\Delta$SfPat resulted in a decrease in the mean diameter of 4.04 mm, demonstrating reduced motility.\\
\begin{figure}
\begin{fullwidth}
\centering
\includegraphics[width=\textwidth]{Figures/Figure 2.pdf}
\caption[Figure 2]{Effects of prophages on biofilm and swimming in \textit{S. fidelis} 3313. (a) Influence of prophages on \textit{in vitro} biofilm formation over 24 hours quantified with crystal violet assay (n=3), (b) role of prophages in swimming quantified as the diameter of spread on soft agar after 24-hours (n=6), and (c) fold-change of \textit{pdeB} (with \textit{Rho} as internal control) from 24-hour biofilm (\textit{in vitro}) (n=4) and 24-hour \textit{in vivo} (n=4). (*= p-value<0.05, **= p-value<0.01; ns= not significant).}
\label{fig:2}
\end{fullwidth}

% Supplementary figure 1
\figsupp[Fig S2]{ qPCR showing relative fold change gene expression with Rho as an endogenous control (a) Cyclic-di-GMP regulators expression in WT and $\Delta$SfPat \textit{in vitro} after 24 hours of exposure (n=4), (b) Cyclic-di-GMP regulators expression in WT and $\Delta$SfPat \textit{in vivo} of \textit{Ciona} MS4 after 24 hours of exposure (n=4). (ns= not significant)}
{\includegraphics[width=\textwidth]{Figures/supp figure 1.pdf}}\label{figsupp:sf1}
\let\figsupp\relax
\end{figure}

\subsection{The influence of a prophage colonization of the \textit{Ciona} gut by \textit{S. fidelis} 3313}

Swimming and biofilm formation behaviors also depend on the modulation of cyclic-di-GMP in the bacteria. Thus, changes in the expression levels of four genes regulating cyclic-di-GMP (\textit{pleD}, \textit{pilZ}, chitinase, and \textit{pdeB}) were measured by qPCR between WT and $\Delta$SfPat. We aimed to identify regulators of the transition from sessile to motile behavior impacted by the deletion of SfPat. No significant change in the expression of the genes in bacteria recovered from 24-hour biofilms (\textit{in vitro}) (\nameref{fig:2} c and Sup fig s1a) was identified. However, when these strain variants were introduced to metamorphic stage 4 (MS4) \textit{Ciona} for 24 hours, the RT-qPCR revealed significant changes in the expression of pdeB (\nameref{fig:2} c and Sup fig s1a). The bacterial gene \textit{pdeB} encodes a phosphodiesterase enzyme that degrades cyclic-di-GMP. By reducing cyclic-di-GMP, pdeB serves as a positive regulator of motility and a negative regulator of biofilm formation \cite{RN73}. The \textit{pdeB} expression was higher in the colonizing WT \textit{S. fidelis} 3313 than the $\Delta$SfPat mutant strain (\nameref{fig:2}c).\\

Swimming and biofilm formation often facilitate bacterial colonization of a host. We investigated whether prophages could impact the ability of \textit{S. fidelis} 3313 to colonize the \textit{Ciona} gut. Colonization assays were performed on MS4 \textit{Ciona} juveniles by exposing animals to  WT or $\Delta$SfPat strains, repeating the experiments six times to account for diverse genetic backgrounds (\nameref{fig:3} a), that is, using gametes from distinct outbred adults. \\

After exposure to the bacterial strains, retention was estimated by recovering bacteria from animals and quantifying colony-forming units (cfu) at different time points. The $\Delta$SfPat strain revealed a statistically insignificant 1.3-to-1.5-fold change in retention compared to the WT strain after 1 hour of bacterial exposure, a time point that mimics initial colonization (\nameref{fig:3} a). However, 24 hours after exposure, WT was over two-fold retained in the gut than the $\Delta$SfPat strain(p<0.05) (\nameref{fig:3} a).\\

To visualize the localization of WT and $\Delta$SfPat mutant strains in the gut, juveniles of MS4 Ciona were exposed for one hour to BacLight Green-stained WT and BacLight Red-stained $\Delta$SfPat strain variants and vice versa (\nameref{fig:3} c-e). The one-hour time point reflects changes in the initial colonization of juveniles.  These experiments revealed a differential localization to the stomach epithelial folds by the WT and $\Delta$SfPat mutant strains. The WT strain typically prefers to occupy the pyloric cecum and the posterior portion of the esophagus and entrance into the stomach (\nameref{fig:3}d and e, Supp Fig. s3a and s3b). Retention of the $\Delta$SfPat mutant was noted at the walls of the stomach during co-exposure (\nameref{fig:3}c).  Retention of the $\Delta$SfPat mutant was observed in the stomach and intestines, and less in the esophagus (Supp fig. s3 c and s3d). These studies suggest spatial and temporal differences in retention by differentially lysogenized strains of \textit{S. fidelis} 3313.\\

\subsection{Host immune discrimination and impact on lysogenized bacteria}

Host immunity also plays an important role in shaping gut homeostasis. Distinct microbes and their antigens and/or metabolites can elicit host immune responses (\cite{RN74}). To determine if the \textit{Ciona} immune system discriminates among \textit{S. fidelis} 3313 strains differing only in the presence or absence of the SfPat prophage, we examined the expression patterns of a secreted immune effector, VCBP-C, among juvenile MS4 during intestinal colonization. Under normal healthy conditions, VCBP-C is expressed and secreted by the gut epithelium and can bind (and opsonize) bacteria within the gut lumen (\cite{RN41}) and influence biofilms \textit{in vitro} (\cite{RN40}). After one hour of exposure to the \textit{S. fidelis} 3313 WT and mutant strains, changes were detected in the expression of VCBP-C. Up-regulation of VCBP-C was observed when juveniles were exposed to $\Delta$SfPat mutant strains of \textit{S. fidelis} 3313, compared to the WT strain (\nameref{fig:4} a) by qPCR(p<0.05). As VCBP-C is a major secreted effector of the gut regulating microbial settlement dynamics, the expression of other innate immune genes was evaluated after 24 hours of exposure to the strains and did not reveal statistically significant responses. However, the lack of statistically significant responses may also be attributed to host genetic diversity. i.e., differing responses to the same strains can obscure signal in transcript pools (\nameref{fig:4} b). \\

Since the presence of SfPat influences host VCBP-C responses, we investigated whether the binding of VCBP-C to bacterial cell surfaces could influence prophage gene expression.  \textit{S. fidelis} 3313 was grown in MB \textit{in vitro} for 24 hours in the presence or absence of 50$\mu$g/ml of recombinant VCBP-C(\cite{RN40}). The supernatant was discarded. RNA was extracted from the biofilms, and gene expression among SfPat open reading frames was monitored, as well as the SOS response regulators, \textit{lexA }and \textit{recA}, in \textit{S. fidelis} 3313. Adherent cultures, or biofilms, were analyzed rather than the supernatant since preliminary experiments revealed that the growth of WT \textit{S.fidelis}  in stationary culrures is influenced by VCBP-C. Interaction of VCBP-C with the WT strain was found to suppress the expression of the structural phage protein P5 of SfPat (\nameref{fig:5}). It was noted that VCBP-C did not significantly alter the expression of \textit{lexA} and \textit{recA}, indicators of the SOS pathway (\nameref{fig:5}). This suggests that VCBP-C binding to the surface of the \textit{S. fidelis} 3313 strain may not influence prophage structural genes via conventional SOS responses. \\

% Main figure
\begin{figure}[ht]
\begin{fullwidth}
\centering
  \includegraphics[width=\textwidth]{Figures/Figure 3.pdf}
  \caption[Figure 3]{The influence of SfPat prophage on gut colonization in \textit{Ciona}. (a)  Results of six biological replicates (n=6, each replicate being a pool of ten juvenile tunicates) of the experimental exposure of \textit{Ciona} MS4 juveniles to either WT or $\Delta$SfPat strains for 1 hr and 24 hrs; retention quantification displayed as a Beeswarm plot of colony-forming units (CFUs). There is significant retention observed in WT after 24 hours. The MS4 juveniles reveal differential colonization of WT and $\Delta$SfPat after one hour of exposure (b-e), where WT strain is stained with BacLight Red and $\Delta$Sfpat is stained with BacLight Green reveal (b) WT is seen localized in the lower esophagus to anterior stomach, while the $\Delta$SfPat deletion strain localized to the hindgut, while (C) the WT is seen localized mostly as a fecal pellet in the center of the stomach while $\Delta$SfPat prefers to localize to the stomach wall. (d) The WT strain is retained in the pyloric cecum. (e) Summary schematic of asymmetric bilaterial views of MS4 animals; top of image is anterior and stomach is posterior. The ventral side is the "En' side, and the dorsal side is the opposite side. The findings can be summarized as such: WT is retained in E and S, in PC, and also in the HG, while the $\Delta$SfPat is retained in the stomach folds, MG, and portions of the HG. Some overlap in signal is noted with yellow coloring. (En = Endostyle, E= Esophagus, S= Stomach, MG= Mid Gut, HG = Hind Gut, PC = Pyloric  Cecum)}
  \label{fig:3}
\end{fullwidth}

% First Supplement
\figsupp[Figure 3—figure supplement 1]
 {WT stained with BacLight Red exposed to \textit{Ciona} MS4 for one hour. (a) WT found in esophagus (E), stomach (S), and fecal pellet (FP). (b) WT also found to occupy the hind gut (HG). (c) WT is retained in the center of the stomach but not the stomach walls.}
{\includegraphics[width=\textwidth]{Figures/supp figure 3.pdf}}

% Second Supplement
\figsupp[Figure 3—figure supplement 2]
{$\Delta$SfPat localization in \textit{Ciona} MS4 after one hour exposure. 
  (a) Stained with BacLight Red, $\Delta$SfPat is found adhered to the stomach (S) and mid gut (MG). 
  (b) Stained with BacLight Green, $\Delta$SfPat is found in the mid gut and hind gut (HG). 
  (c) $\Delta$SfPat is more localized in the stomach fold than the center of the stomach, with presence in the mid gut.}
  {\includegraphics[width=0.8\textwidth]{Figures/supp figure 4.pdf}}

\end{figure}
 
\begin{figure}[ht]
\begin{fullwidth}
  \includegraphics[width=\textwidth]{Figures/Figure 4.pdf}
  \caption [Figure 4]{The influence of prophages on host gene expression. (a) VCBP-C gene expression in MS4 juveniles after one hour of exposure to \textit{S. fidelis} 3313 strains (n=4), (b) Survey of additional innate immune gene expression in MS4 juveniles after 24-hour exposure to WT or $\Delta$SfPat mutant strains (n=3). Actin is the internal control. (*= p-value<0.05, **= p-value<0.01, ns = not significant).}
  \label{fig:4}
\end{fullwidth} 
\end{figure}
\begin{figure}[h]

\begin{fullwidth}
  \includegraphics[width=\textwidth]{Figures/Figure 5.pdf}
  \caption [Figure 5]{Lysogen gene expression in response to host immune effector binding. Gene expression of SfPat structural protein p5, \textit{recA} and \textit{lexA} of WT strain grown as a 24-hour biofilm while exposed to 50 $\mu$g/ml VCBP-C. Rho is the internal control (n=4). (*= p-value<0.05, ns = not significant).}
  \label{fig:5}
 \end{fullwidth}  

\figsupp[Figure 5 supplement]
{Rho is the most stable gene across strains when tested \textit{in vitro} (n=3)}
{\begin{fullwidth}
\includegraphics[width=1.2\textwidth]{Figures/supp figure 2.pdf}
\end{fullwidth}}

\end{figure}

\section{Discussion}

In this report, we utilized a phage-deletion strategy to study the influence of a prophage (SfPat) on a gut symbiont (\textit{S. fidelis} 3313) of the model invertebrate, \textit{Ciona robusta}. Using RAST annotations and deduced amino acid BLAST (BLASTx), we propose that the SfPat is a distant relative of the PM2 prophage (Supplementary Table 1, \cite {RN121}).  Deletion of the PM2-like SfPat prophage from \textit{S. fidelis} 3313 revealed phage-mediated influence on colonization dynamics. We find that the deletion of the PM2-like prophage decreases swimming behaviors while increasing biofilm formation. These bacterial phenotypes influence host immune responses in ways that may influence differential retention in the gut if \textit{Ciona}. Our identification of a prophage that interferes with biofilm formation in a \textit{Shewanella} strain contrasts with reports demonstrating an increase in biofilms resulting from the incorporation of exogenous DNA following cell lysis (\cite{RN42, RN43}). Our study is the first to implicate prophage-induced change in motility for \textit{Shewanella} spp, and it supports the reasoning that prophages can impact their hosts by modulating traits via diverse mechanisms. An influence on motility has also been linked to a prophage in \textit{Pseudomonas aeruginosa} (\cite{RN78}). \\

SfPat in \textit{S. fidelis} 3313 appears to have a variable influence on bacterial retention in the gut of \textit{Ciona}. As feeding is initiated in MS4 \textit{Ciona} juveniles, food (or bacteria in the environment, natural or artificially introduced) accumulates in the gastrointestinal tract and on average takes about 45 minutes to begin exiting the anus and atrial siphon as fecal pellets.  Thus, we monitored and compared transit and retention of introduced bacteria in the gut of MS4 juveniles at one and 24 hrs after introduction. SfPat deletion reveals a prophage-mediated influence on gut retention and localization. For example, within one hour of exposure, fecal pellets begin to form in the stomach that are enriched for the WT \textit{S. fidelis} 3313; however, the WT strain then appears to be retained in the posterior portion of the esophagus (just before the entrance into the stomach) as well as in the pyloric cecum (which is a small outpouching just ventral and posterior to the stomach). Instead, the $\Delta$SfPat mutant strain appears to be retained in the stomach folds and the intestines, and not the esophagus.  It should be noted that the base of the stomach folds is where secretion of VCBP-C occurs \cite{RN40, RN41, RN115}. The overall retention of the two strains did not vary significantly within the first hour of colonization. However, after 24hrs, the retention of the WT strain was significantly greater than that of $\Delta$SfPat despite the continued feeding behavior of the MS4 juveniles. The increase in pdeB activity of the WT \textit{in vivo} (\nameref{fig:2}) might indicate that the secondary messenger, cyclic-di-GMP, may be reduced in the WT, possibly leading to increased biofilm formation and reduced motility(\cite{RN75}). Thus, the presence of SfPat influences niche preference and retention. \\

In addition to host genetics obscuring the influence of prophages on colonization of the gut, other biophysical factors that include host immune effectors play crucial and often silent roles in influencing bacterial settlement dynamics. For example, human secretory immunoglobulin A (SIgA) has been shown to enhance and often favor settlement of bacteria both \textit{in vitro} and \textit{in vivo} (\cite{RN80,RN81,RN82,RN83}), raising a basic question as to whether this phenomenon is more widespread among other secretory immune effectors present in mucosal environments of animals (\cite{RN84}). We speculate that while prophages likely impact the behavior of lysogenized bacteria in ways that can influence colonization dynamics, interaction with VCBP-C on the mucosal surface of the \textit{Ciona} gut likely further influences settlement behaviors (\cite{RN40,RN84}). Importantly, we show here that the influence of the SfPat prophage on bacterial physiology (the WT strains) leads to a reduced expression of \textit{Ciona} VCBP-C in the first hours of bacterial retention in the gut (an indicator that the host has detected the exposure). It remains to be shown if prophages stimulate the production of a bacterial metabolite with immunomodulatory properties or if the host immune system responds to differences in bacterial behaviors or traits, as suggested in the $\Delta$SfPat deletion mutant. \\

Metagenomic sequencing of gut microbes from healthy humans has revealed that temperate lifestyles are prevalent among phages from these ecosystems (\cite{RN85,RN86, RN87}), an observation also made in the \textit{Ciona} gut (\cite{RN15}). Various environmental triggers, such as UV light and mutagenic agents like mitomycin C, have been shown to induce a switch from the temperate to lytic cycle via the SOS response, a cell-wide response to DNA damage that can promote survival (\cite{RN88}). Since, VCBP-C is an immune molecule in the gut that can interact with bacteria, it could influence prophage induction. However, we find that VCBP-C binding on the surface of WT \textit{S. fidelis} 3313 leads to a reduction in the expression of an important SfPat structural protein P5, suggesting a limitation in SfPat induction in the presence of VCBP-C. No significant changes in lexA/recA expression were observed upon VCBP-C exposure/binding, suggesting a lack of SOS response when exposed to this immune effector. The various mechanisms by which prophages shape colonization behaviors among gut bacteria of animals remains unclear. While the data reported here are only based on one bacterial strain that colonizes the \textit{Ciona} gut, we find that WT \textit{S. fidelis} colonizes the gut with reduced activation of VCBP-C gene expression compared to $\Delta$SfPat, a trait that may be important in shaping colonization outcomes. We speculate that these observations are more widely applicable since lysogens are so abundant in animal microbiomes. Under normal conditions, VCBP-C protein is present in copious amounts and tethered to chitin-rich mucus lining the gut, as revealed by immunohistochemical staining (\cite{RN40}). Therefore, overexpression of VCBP-C is not necessarily helpful, and can correspond to the induction of additional inflammatory responses, including an overproduction of mucus. Thus, regulation of the production of additional VCBP-C likely serves important roles in influencing colonization dynamics.\\

Since colonization of animal mucosal surfaces is an ancient process (\cite{RN32}), prophages and their integration into bacterial genomes have likely evolved to provide fitness benefits in often challenging environments like the gut lumen. Determining the role of animal immunity and prophages in these exchanges is of broad interest. Immune effectors like VCBPs, which undoubtedly possess broad specificities, can bind a range of bacterial hosts; however, it remains to be shown if they bind lysogenized bacteria with different affinities than their prophage-free counterparts. Prophages can also be induced to generate lytic particles that can influence gut microbiome structure, serving as an indirect form of protection for the host \cite{RN25,RN89}. Prophages can also contribute to the transfer of virulence factors (\cite{RN24,RN90}). Retention of lysogens may be preferred if the prophages provide competitive fitness and retention in the gut. Since lysogens are integral in animal development, immunity, and metabolism(\cite{RN120}), there is a tripartite interplay required for survival, a snapshot of which is shown here.\\

\section{Methods and Materials}

\subsection{Culture and growth conditions}

\textit{S. fidelis} 3313 used in this study was originally isolated from the gut of \textit{Ciona robusta} obtained from Mission Bay, CA, USA, as previously described \cite{RN33}. The bacterium was cultured using Difco marine agar 2216 (MA) (Fisher Scientific, Hampton, NH) and marine broth (MB) at room temperature (22-24 $^{\circ}$C). Subsequent genetic manipulations were performed on strains grown in LB/MB, which consists of a mixture of 75\% LB (Lysogeny Broth (Luria), Fisher Scientific, Hampton, NH) and 25\% MB. Strains are listed in \nameref{tab:strainslist}.\\
\begin{table} [H]
\caption[Table 1]{\label{tab:strainslist}All \textit{S. fidelis} 3313 strains are submitted under the BioProject \href{https://www.ncbi.nlm.nih.gov/bioproject/903273}{PRJNA 90327} on NCBI, Accession:  SAMN31793880 ID:31793880}
\begin{tabular}{l l l}
\toprule
\textbf{Organisms}&\textbf{Phenotype}&\textbf{NCBI RefSeq Assembly}\\
\midrule
JG4066&WT&GCF\_033441085.1\\
JG3862&$\Delta$SfPat&GCF\_033441065.1\\
\bottomrule
\end{tabular}
\end{table}

\subsection{Prophage deletion}

SfPat was targeted for deletion from \textit{S. fidelis} 3313 using homologous recombination methods adapted from \cite{RN51} to produce knockout mutant strains. First, a pSMV3 suicide vector \cite{RN51} was designed with ~700 bp regions corresponding to the upstream and downstream sequence of the prophage ( \nameref{tab:deletelist}). These flanking regions were amplified and ligated using overlap extension PCR, then directionally inserted into the vector with the restriction enzymes BamHI and SacI (\nameref{tab:plasmidslist}) \cite{RN52}. Plasmid conjugation was then performed by inoculating a colony of \textit{S. fidelis} 3313 into a culture of \textit{E. coli} containing the desired suicide vector on an LB/MB agar plate for two hours followed by primary selection after 24 hours at RT on LB/MB + 100  $\mu$g/ml Kanamycin plates and a counter selection on LM/MB + 10$\%$  Sucrose at RT. Merodiploids and deletion mutants were verified by PCR.
Illumina sequencing (MiGS, University of Pittsburgh) confirmed the deletion of SfPat and the evaluation of any additional genetic changes or mutations (Figure s1a). \\

\begin{table}[H]
\caption[Table 2]{\label{tab:deletelist}Primer sequences on \textit{S. flidelis} 3313 used for generating $\Delta$ SfPat deletion suicide vector and for deletion verification}
\begin{tabular}{l l}
\toprule
\textbf{Primer ID} & \textbf{Sequence (5’-3’)} \\
\midrule
EDK77 & AAATGGATCCCGATCAGCCTGCTAGTTTATT \\
EDK78 & ACGGAATAGGTTGAATGCGACTCAGGC \\
EDK79 & TCGCATTCAACCTATTCCGTCATGTTTAGCC \\
EDK80 & ACATGAGCTCGATGCAGATAAAGAGCCGTAAA \\
EDK81 & GTTTATTTTGTGGCAATCGCA \\
EDK82 & GGTAGCAGTGCTTAAACGAT \\
\bottomrule
\end{tabular}
\end{table}

\begin{table}[ht]
\caption[Table 3]{\label{tab:plasmidslist}Plasmids and strains used in the study}
\begin{tabularx}{\textwidth}{X X X}
\toprule
\textbf{Plasmids/ Strains} & \textbf{Genotype} & \textbf{Source/ Reference} \\
\midrule
\textit{E. coli} UQ950 & \textit{E. coli} DH5$\alpha$ \textnormal{$\lambda$(pir) host for cloning; F$^{-}$ $\Delta$ (argF$^{-}$ac)169 $\Phi$ 80dlacZ58($\Delta$ M15) glnV44(AS) rfbD1 gyrA96(NalR) recA1 endA1 spoT1 thi$^{-}$1 hsdR17 deoR $\Lambda$ pir $^{+}$} & \cite{RN51} \\
pSMV3$-\Delta$ SfPat & pSMV3 with 778 bp upstream and 779 bp downstream of flanking regions of SfPat & This study \\
\bottomrule
\end{tabularx}
\end{table}

\subsection{\textit{S. fidelis} crystal violet biofilm assay}

WT and $\Delta$SfPat strains were cultured in MB overnight at RT and then diluted to $10^7$ cfu/ml in MB. Cultures were brought to a final volume of 2 ml of MB in 12-well dishes and incubated at RT for 24 hours to examine biofilm development. Each variable was tested in technical duplicate. The biofilms were quantified by crystal violet staining as previously described \cite{RN56}. Briefly, the supernatants were aspirated after 24 hours of incubation and the biofilms were dried and stained with 0.1\% crystal violet for 10 min. The stained biofilms were then gently washed with deionized water and the amount of biofilm produced was quantified as the intensity of the stain $(OD_{570})$ after the biofilm-bound crystal violet was extracted from the biofilm with 30\% acetic acid. All biofilm assays were performed at least in triplicates. \\

\subsection{Motility assay}

Soft-agar overlay motility assays were carried out in 12-well dishes to compare swimming behaviors \cite{RN57, RN58}. Briefly, a single colony from an overnight streaked LB/MB plate was picked using a sterile tooth pick and then stabbed onto the center of soft agar (containing LB/MB and 0.5\% low-melt agarose) and incubated at RT overnight. The results were recorded as the distance traveled (in millimeters) by the bacteria from the inoculation zone. Each variable was tested in duplicate. Two perpendicular distances from the inoculation zone were recorded for each technical replicate and averaged for each well.\\

\subsection{\textit{Ciona} mariculture} 

The \textit{in vivo} colonization experiments were performed on animals reared under conditions termed “semi-germ-free” (SGF), which include minimal exposure to marine microbes. SGF conditions include animals harvested under conventional approaches \cite{RN59} but permanently maintained in 0.22 $\mu$m-filtered, conditioned artificial seawater (cASW), handled with gloves, and lids only carefully removed for water/media changes. cASW is prepared by conditioning ASW made with Instant Ocean$^\text{TM}$ in an in-house sump-aquarium system containing live rock, growth lights, and sediment from San Diego, California; salinity is maintained at 32-34 parts per thousand (or grams per liter). Compared to germ-free \cite{RN60} or SGF, conventionally-reared (CR) includes a step-up exposure to 0.7$\mu$m-filtered cASW that increases exposure to marine bacteria during development. The SGF approach is considered an intermediate method of rearing that includes minimal exposures to microbial signals during development (unpublished observations). The animals were reared at 20 $^{\circ}$C from larval to juvenile stages. The \textit{Ciona} were collected from Mission Bay, California in order to produce juvenile organisms for each biological replicate. These wild-harvested animals provide a wider genetic diversity compared to traditional model systems, where genetic diversity has been reduced or eliminated through controlled breeding practices. \\

\subsection{Gut colonization assays}

Both bacterial strains were grown overnight at RT in MB and diluted to $10^7$ cfu/ml in cASW after repeated washes. Metamorphic stage 4 (MS4) animals reared in six-well dishes in cASW were exposed to 5 ml of $10^7$ cfu/ml bacteria in each well for one hour or 24 hours. Co-exposure studies were also performed, where 2.5 mL (or 1:1) of each culture prepared above was mixed to form a total volume of 5 ml.  MS4 animals are considered part of the 1st ascidian stages (post-settlement stages 1-6, whereas stages 7-8 and onwards are 2nd ascidian stages and reflect young adult animals). MS4 juveniles can be identified as having a pair of protostigmata, or gill slits, on each side of the animal (\cite{RN34}). These juveniles first initiate feeding via newly developed and opened siphons; before this, the gut remains closed, and the interior lumen is unexposed to the outside world. Following this initial exposure or colonization, for various time intervals, the plates were rinsed multiple times with cASW and replaced with fresh cASW. Ten juveniles were chosen randomly for each treatment, pooled, and homogenized with a plastic pestle; live bacteria were counted by performing serial dilutions and enumerating colony-forming units (cfu) via spot-plating assays (\cite{RN61, RN62}).  Each graphed data represents a biological replicate dataset from genetically distinct/ diverse backgrounds of \textit{Ciona} (represented by separate live animal collection and spawning events). Statistical significance was calculated using the Wilcoxon t-test by pooling data across six genetically diverse biological replicates. \\

Live bacteria in the gut were visualized using BacLight stains and previously described fluorescently labeled bacteria \cite{RN6}. For BacLight staining, 1 ml of bacterial cultures were grown overnight at RT, pelleted, washed twice with cASW, and stained with 4 $\mu$l of BacLight Red (Invitrogen Cat no B35001) or BacLight Green (Invitrogen Cat no B35000) for 15 mins in the dark. The cultures were stained with alternate dyes in different replicates to get unbiased data from changes in fluorescence. The stained cultures were washed twice with cASW, and then diluted to $10^7$ cfu/ml with cASW. MS4 animals were grown in 6-well dishes were then exposed to 5 ml of this culture. Bacteria in the gut of animals were visualized after one hour on a Leica DMI 6000B stereoscope with a CY5 fluorescent filter for BacLight Red and GFP filter for BacLight Green; and imaged and captured with a Hamamatsu ORCAII camera (model C10600-10B-H) and processed with the MetaMorph 7.10.4 imaging suite (Molecular Devices, Downingtown, PA). 

\subsection{Differential transcript level studies}

To determine if the \textit{Ciona} innate immune system can recognize and respond to unique mutant strains, which differ only in the presence or absence of prophages, candidate immune response markers were examined using reverse transcription quantitative PCR (RT-qPCR). RNA  was extracted using the RNA XS kit (Macherey-Nagel, Cat no 740902) from MS4 \textit{Ciona} juveniles exposed to either the WT or $\Delta$ SfPat strain. Complementary DNA (cDNA) synthesis was performed with oligo-dT primers and random hexamers using the First Strand cDNA Synthesis Kit (Promega Cat no A5000) following the manufacturer’s instructions. The amplification was set with the qPCR kit (Promega Cat no A6000) and carried out on an ABI7500 with an initial melting temperature of 95$^{\circ}$C for 2 mins and 40 cycles of 95$^{\circ}$C for 15 sec and 60$^{\circ}$C for 1 min. The innate immune genes examined and their primers are reported in Table \ref{tab:cionaprimerlist}. Results from four distinct biological replicates are presented. Each replicate includes pooled \textit{Ciona} juveniles from at least two wells of a 6-well dish. \textit{Ciona} actin was referenced as an endogenous control. Data was analyzed using $\Delta\Delta$Ct method \cite{RN64} and the ABI7500 software suite. For $\Delta\Delta$Ct method, the Ct values were first normalized to an endogenous control gene and further normalized to the reference samples, here the WT strain.\\

To understand bacterial genes that are differentially regulated due to the presence or absence of prophages, transcript levels were studied \textit{in vitro} and \textit{in vivo}.  The bacterial strains were grown in six-well dishes using the same methodology described for biofilm assays. To understand if the host immune molecule VCBP-C induced prophages, WT was cultured in six-well dishes as described in the biofilm section in the presence or absence of 50  $\mu$g/ml VCBP-C in Marine Broth \cite{RN40}. After 24 hours, the supernatant was discarded for both experiments, and RNA from the biofilm was extracted using the Zymo Research Direct-zol kit. cDNA synthesis was performed using random hexamers as primers, and qPCR was conducted as described above. The targeted bacterial genes are described in (Table \ref{tab:bacprimerlist}). Different housekeeping genes were examined for the different treatments, and the Ct values were used to identify the most stable reference gene to be used as an endogenous control. Rho was identified as the most stable reference gene using RefFinder, which utilizes Bestkeeper, GeNorm, Normfinder, and comparative $\Delta \Delta$Ct methods  (\nameref{fig:5} supplement 1) \cite{RN99, RN64, RN100, RN92}. \\

\begin{table}[h]
  \centering
  \scriptsize
  \caption[Table 4]{\label{tab:cionaprimerlist}\textit{Ciona} genes targeted and the necessary reverse transcription-qPCR primers}
  \begin{tabularx}{\textwidth}{>{\raggedright\arraybackslash}p{1.8cm} >{\raggedright\arraybackslash}p{3 cm} >{\raggedright\arraybackslash}p{2.5cm} >{\raggedright\arraybackslash}p{3.5cm} >{\raggedright\arraybackslash}p{1.9cm}}
    \hline
    \textbf{Gene} & \textbf{Function} & \textbf{Genbank accession no.} & \textbf{Primer (5’-3’)} & \textbf{Reference} \\ \hline
    VCBP-C-Fwd & Secreted immune effector in the gut & HQ324151 & AGACCAACGCCAACACAGTA & \cite{RN91} \\
    VCBP-C-Rev & & & CCCCATACATTGCAGCATTTC & \\ [1ex]
    
    Actin–Fwd & Cytoskeletal actin  & AJ297725 & CCCAAATCATGTTCGAAACC & \cite{RN91} \\
    Actin-Rev & \textbf{Reference gene}& & ACACCATCACCACTGTCGAA & \\[1ex]
    
    IL17-1-Fwd & Interleukin 17 & NM\_001129875.1 & AGGTTAAGAATCCCTATGGTGC & \cite{RN110} \\
    IL17-1-Rev & Effector cytokine& & CAAAGGCACAGACGCAAAGG & \\[1ex]
    
    IL17-1R-Fwd & Interleukin 17 Receptor & NM\_001245045 & TGTTGGCATGAGTGTTCGGT & \\
    IL17-1R-Rev & & & AGTTGGTTCTGCCCCAAAGT & \\[1ex]
    
    NF$\kappa$-B-Fwd &Immune regulatory & NM\_001078304 & TGTCGCTTGTCGTCATGGAA & \cite{RN110}\\
    NF$\kappa$-B-Rev &Transcription factor & & AACACCCAAGACCGTCGAAA & \\[1ex]
    
    TNF$\alpha$-Fwd & Tumor Necrosis Factor & NM\_001128107 & TTCAGAAAGATTGGACGACGA & \cite{RN110} \\
    TNF$\alpha$-Rev &Inflammatory effector & & TCGTTTAGAAATGCTGCTGTGG & \\[1ex]
    
    C3A-R-Fwd & Complement C3& NM\_001078552 & TTGTAAGCTGGCACAAGGTGT & This study \\
    C3A-R-Rev & Inflammatory mediator& & GACCGTAGTCTGGTAGAGGTC & \\[1ex]
    
    MBL-Fwd & Mannose Binding Lectin & NM\_001167707.2 & TTATTGATGGGAAAGTTTGGT & This study \\
    MBL-Rev & & & TAACATCTCTGTTCTTGGGTC & \\
    \hline
  \end{tabularx}
\end{table}
\begin{table}[h]
  \centering
  \scriptsize % Further reduce font size to ensure the table fits
  \caption[Table 5]{\label{tab:bacprimerlist}\textit{S. fidelis} 3313 genes targeted and the necessary reverse transcription-qPCR primers}
  \begin{tabularx}{\textwidth}{>{\raggedright\arraybackslash}p{1.8cm} >{\raggedright\arraybackslash}p{2.8cm} >{\raggedright\arraybackslash}p{2.5cm} >{\raggedright\arraybackslash}p{4cm} >{\raggedright\arraybackslash}p{1.9cm}}
    \hline
    \textbf{Gene} & \textbf{Function} & \textbf{Genbank accession no.} & \textbf{Primer (5'-3')} & \textbf{Reference} \\ \hline
    pleD-Fwd & Regulates production of cyclic di-GMP & NZ\_KI912459.1 & CTCTTCACCGCCACTTCTT & This study \\
    pleD-Rev & & & GGTGTGGTCTCTTATGCCTATC & \\[1ex]
    Chitinase-Fwd & Chitin utilization gene & NF027718.3 & CAGTGTAGCTAAGTCGTCATC & This study \\
    Chitinase-Rev & & & CGCCAACCAGTGCTTTATTG & \\[1ex]
    pilZ-Fwd & Type IV Pilus control protein & NZ\_JADX01000014.1 & TGGCAAGGTCGTTTGGATTA & This study \\
    pilZ-Rev & & & AGGCAAGCTCACTGGAAAG & \\[1ex]
    pdeB-Fwd & Phosphodiesterase & NF012772.3 & GCATCAGGGCTCTTACCAATAG & This study \\
    pdeB-Rev & & & GAGGCGGTGATCCTTACAGATA & \\[1ex]
    recA-Fwd & Bacterial recombinase / Bacterial reference gene& NZ\_KI912459.1 & CGTAGTGGTGCGGTAGATGT & This study \\
    recA-Rev & & & CGCATTGCTTGGCTCATCAT & \\[1ex]
    lexA-Fwd & Regulator of recombinase & NZ\_JADX01000011.1 & TGACCCAGCTATGTTCCGCC & This study \\
    lexA-Rev & & & GCTCAACCTTGTGTACGGCG & \\[1ex]
    Rho-Fwd & \textbf{Bacterial reference gene} & NZ\_JADX01000026.1 & CACGTACAAGTTGCCGAAATG & This study \\
    Rho-Rev & & & CAAGACGGGTGATAGAGTCAAG & \\
    gyrB-Fwd & \textbf{Bacterial reference gene} & AM229309.2 & TCACGAGCATCATCACCCGT & This study \\
    gyrB-Rev & & & GGCTTCCGTGGTGCGTTAAC & \\
    \hline
  \end{tabularx}
\end{table}

\subsection{Statistical analysis and data visualization}

Statistical analysis and data visualization were carried out in R, version 4.2 \cite{RN65}. Data were plotted with ggplot 3.3.5 \cite{RN69}; the Beeswarm plot was constructed using ggbeeswarm 0.6 \cite{RN67}. Beeswarm plots and statistics for motility assays were calculated using replicate averages \cite{RN68}. Statistical significance was calculated using ggsignif package 0.6.3 or ggpubr0.4.0 \cite{RN69,RN70}. If the data were found to be normally distributed by Shapiro’s test, then significance was calculated using an unpaired t-test. The Wilcoxon signed-rank test was used to calculate the significance of non-parametric data.

\subsection{Author contributions}

O.N. designed, executed, and analyzed experiments and wrote and edited the manuscript. S.L.G., N.P., C.G.F.A., F.N, A.L., and E.D.K performed experiments and provided feedback and approved the manuscript. M.N.Y., S.J.L., and B.A.L. performed genome sequence analysis, assembly, and provided feedback and approved the manuscript. M.B. and J.A.G. helped interpret data, provided feedback, and approved the manuscript. L.J.D. helped design experiments, analyze data, and helped write, edit, and approve the manuscript.

\subsection{Funding}

This project was supported by NSF IOS-1456301 (L.J.D. and M.B.), NSF MCB-1817308 (L.J.D.), NSF IOS-2226050/51 (L.J.D. and J.G.), internal awards from the MCOM Deans Office, Ann and Andrew Hines Endowed Chair in Molecular Genetics, and the USF Institute for Microbiomes to L.J.D. Additional funding included an NSF GRFP award to B.A.L., and a New Investigator Award to O.N. by USF.

\section{Acknowledgments}

The authors acknowledge the expertise of Gary W. Litman and John P. Cannon for expert feedback and guidance on earlier efforts of the project, and anonymous reviewers who helped improve earlier versions of the manuscript.
\bibliography{elife-sample}

%%%%%%%%%%%%%%%%%%%%%%%%%%%%%%%%%%%%%%%%%%%%%%%%%%%%%%%%%%%%
%%% APPENDICES
%%%%%%%%%%%%%%%%%%%%%%%%%%%%%%%%%%%%%%%%%%%%%%%%%%%%%%%%%%%%

\appendix

\end{document}
