## Supplementary figures and images for "Prophage regulation of *Shewanella fidelis* 3313 motility and biofilm formation: implications for gut colonization dynamics in *Ciona robusta*"

### Figure 1.pdf

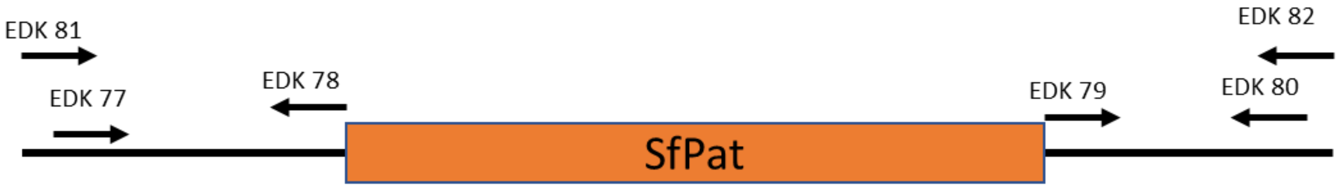

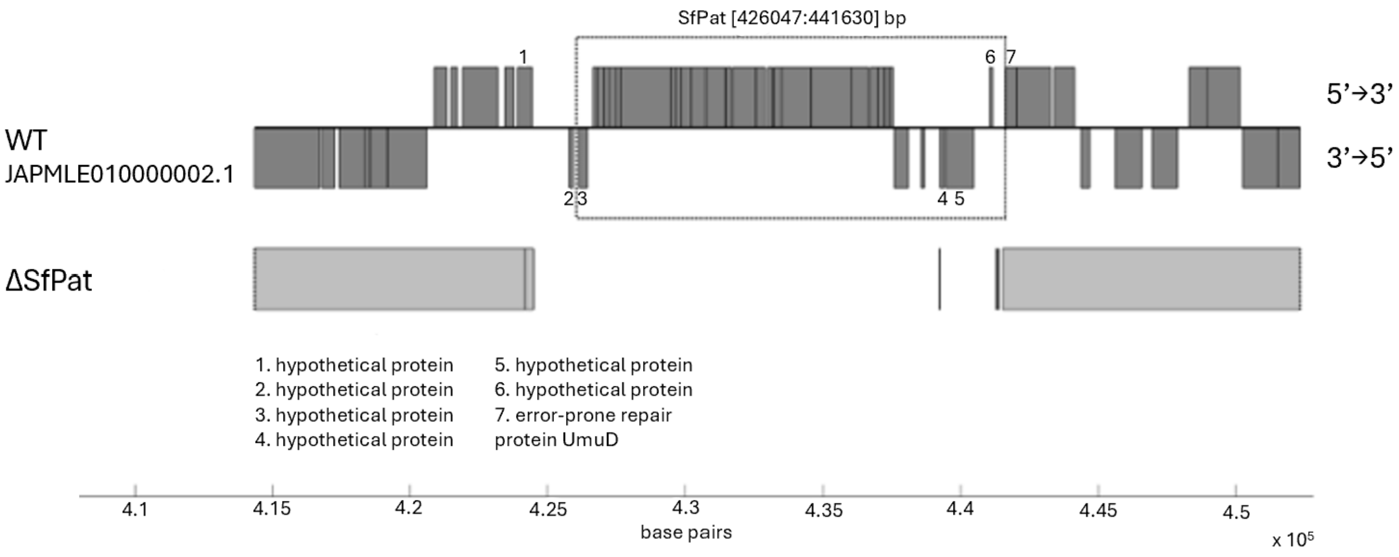

### Figure 2.pdf

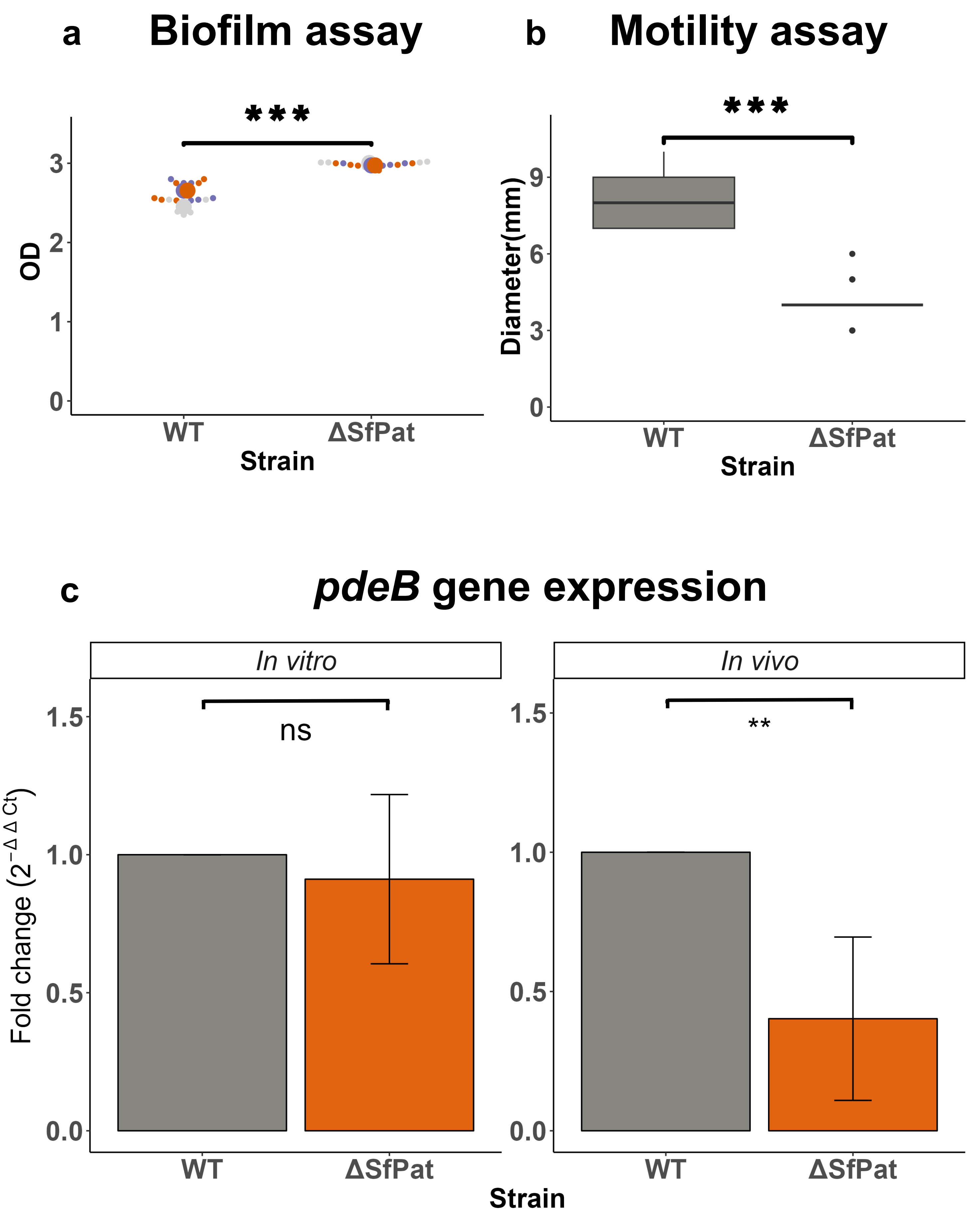

### Figure 3.pdf

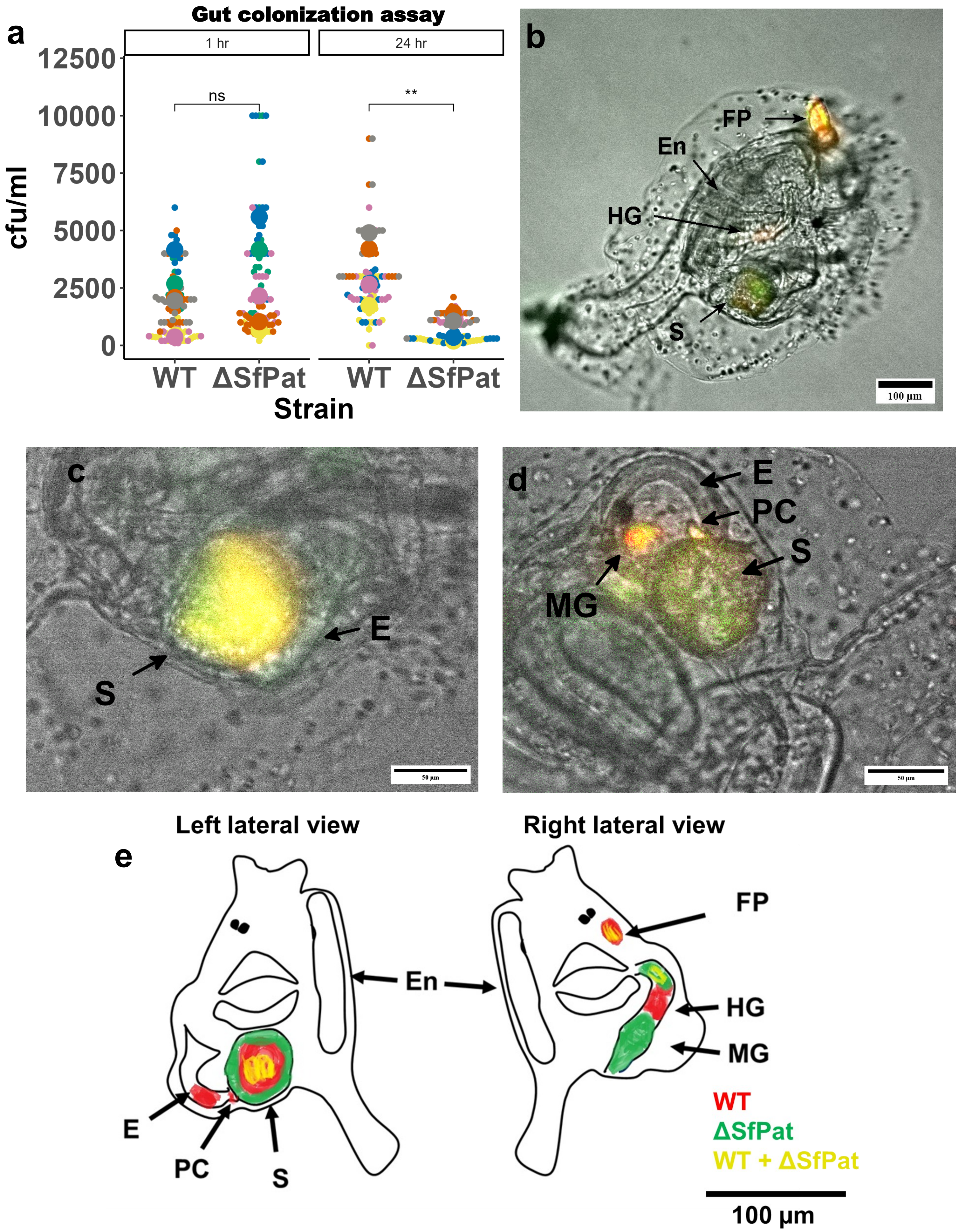

### Figure 4.pdf

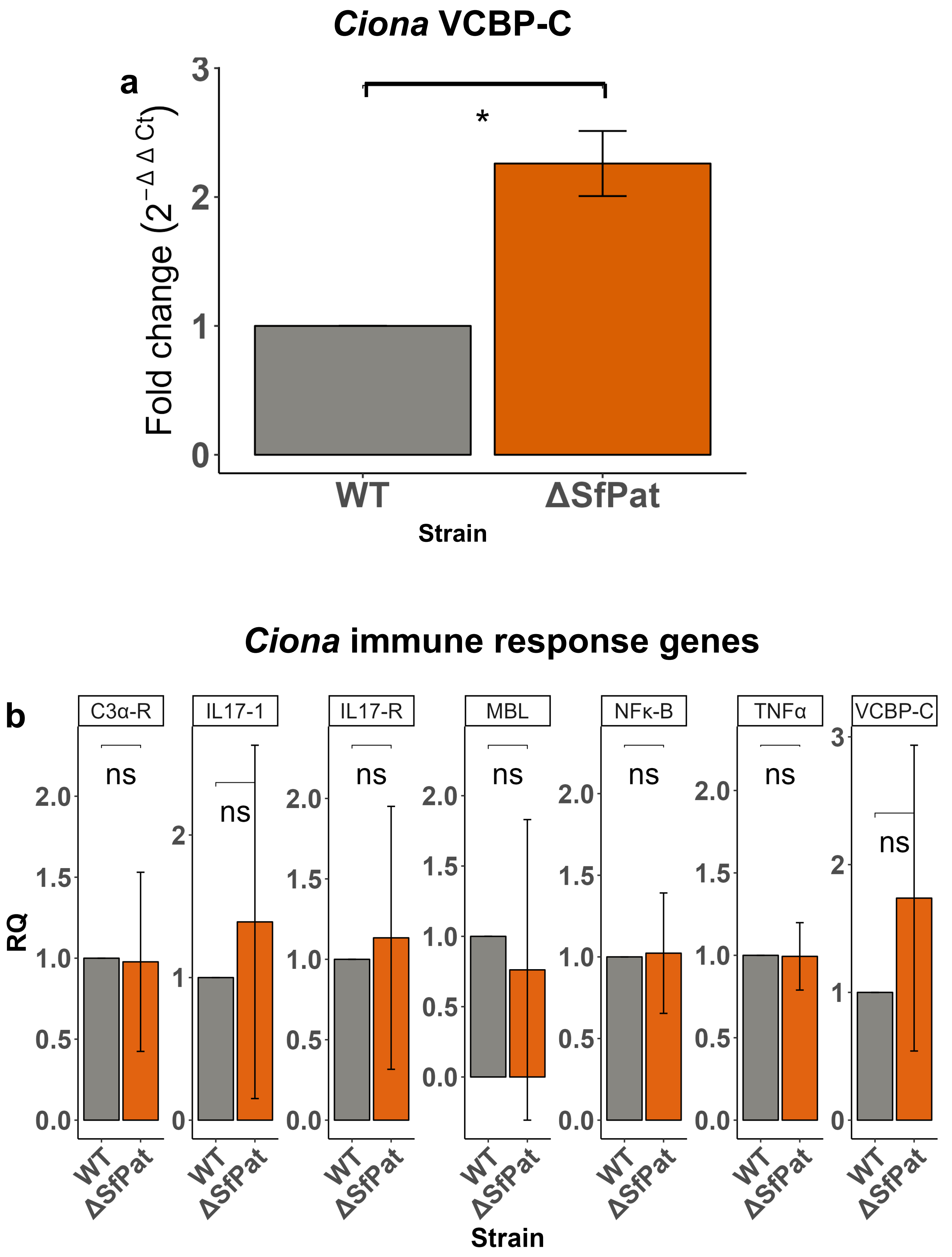

### Figure 5.pdf

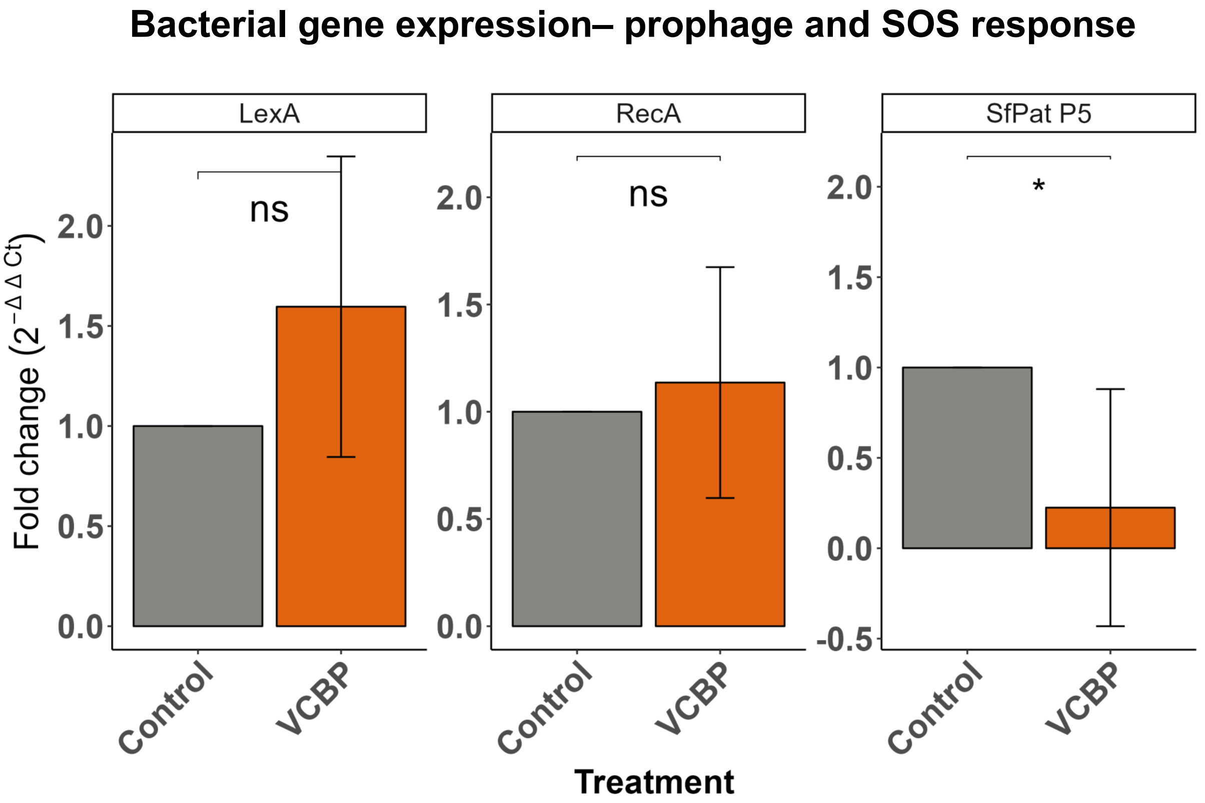

### supp figure 1.pdf

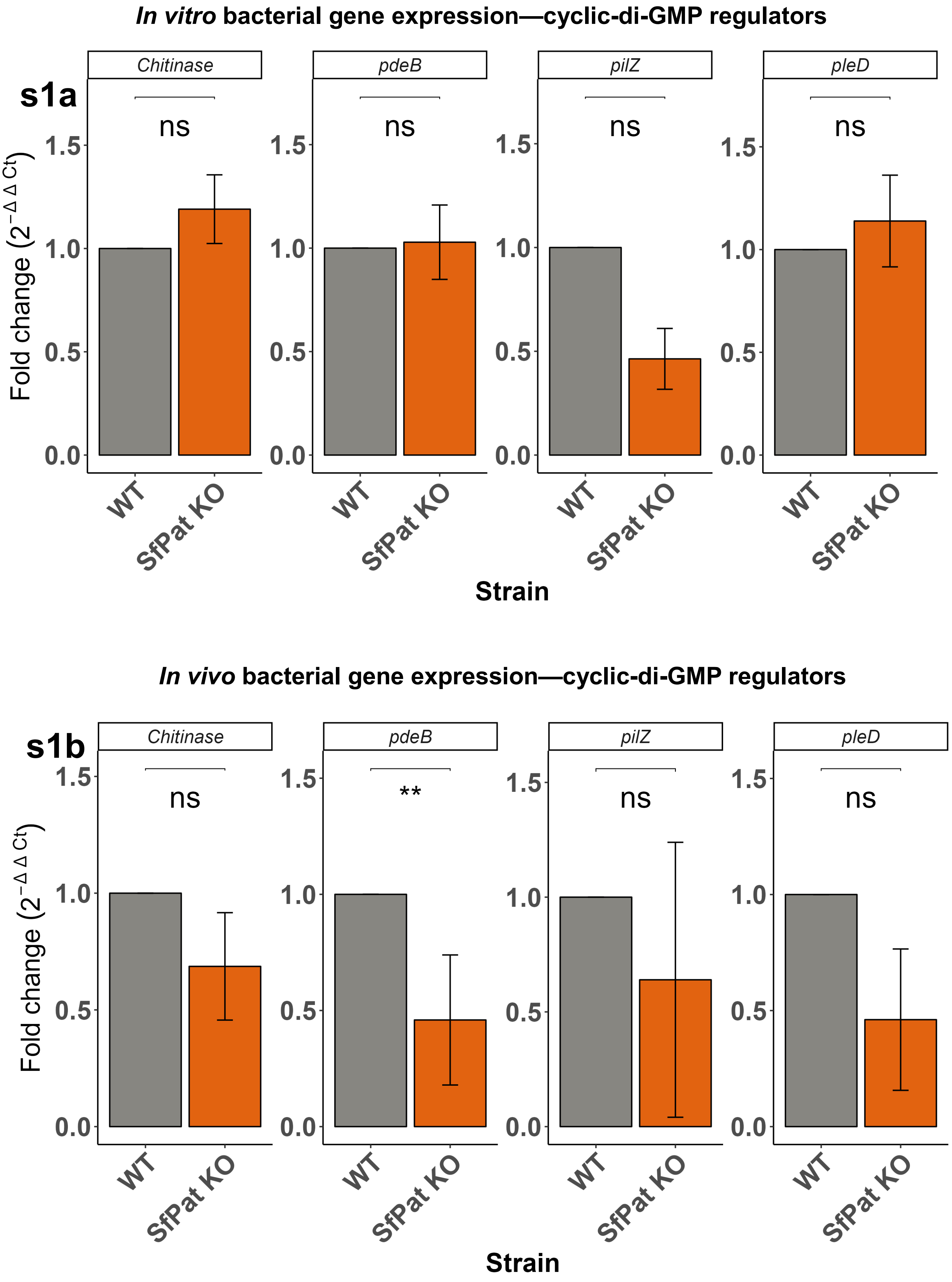

### supp figure 2.pdf

| Genes | Geomean of ranking values |
| --- | --- |
| rho | 1.00 |
| RecA | 1.86 |
| gyrB | 2.71 |

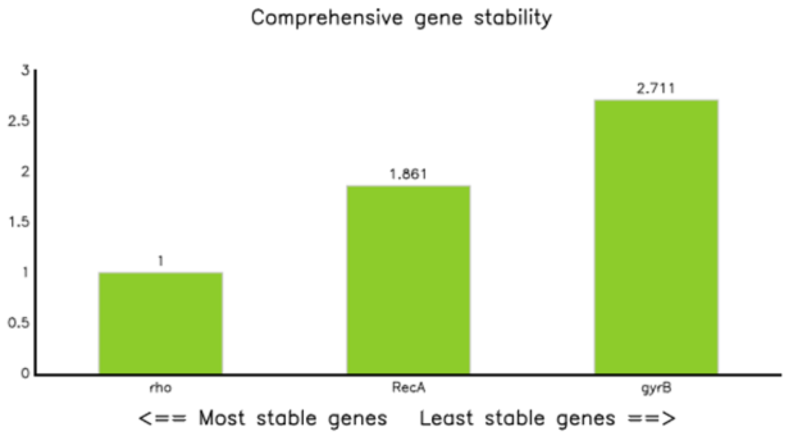

### supp figure 3.pdf

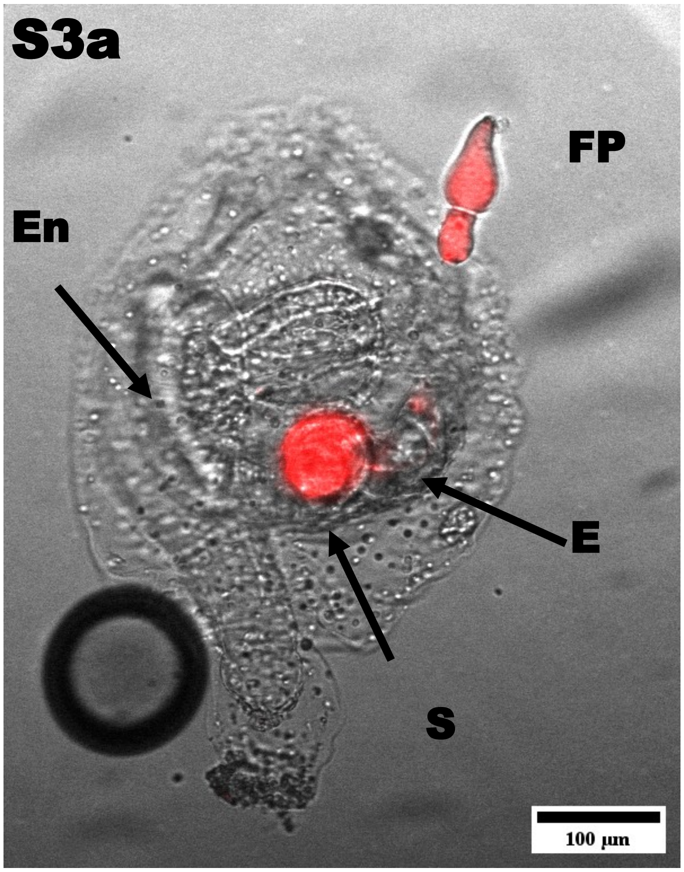


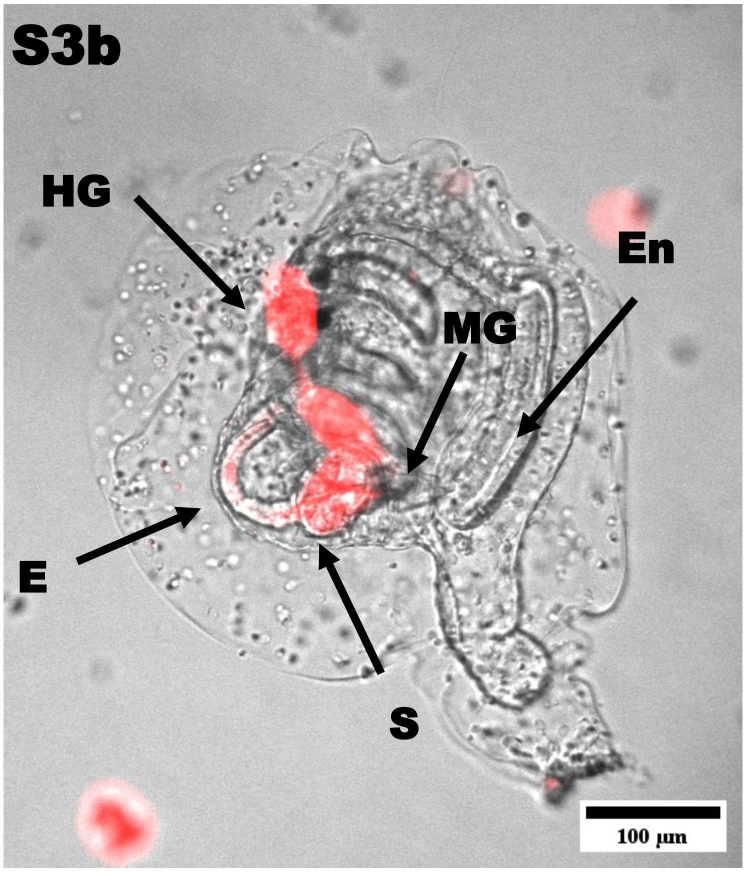


S3c

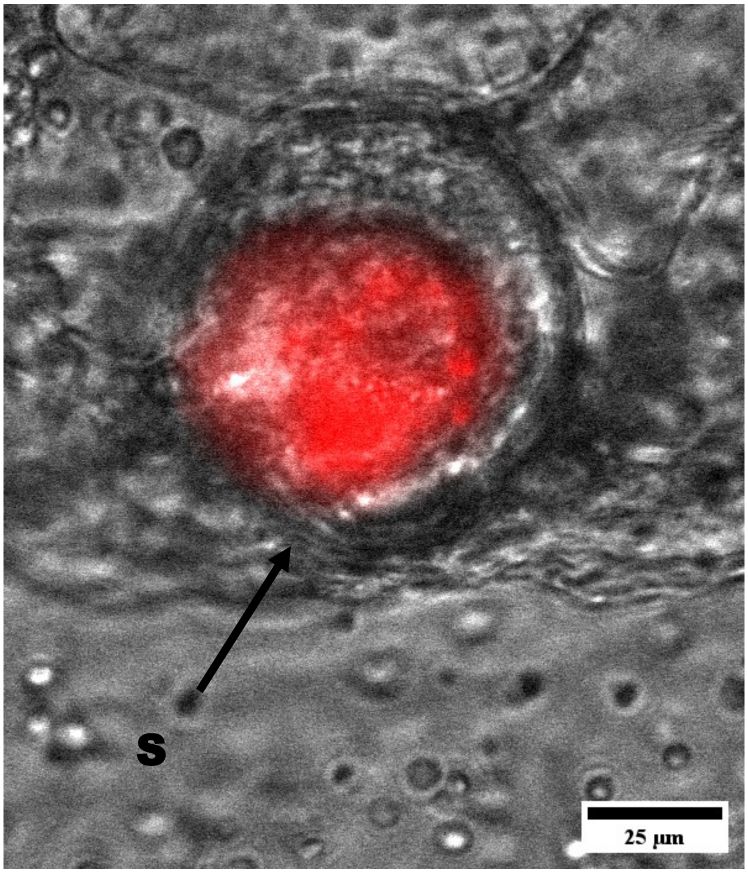

### supp figure 4.pdf

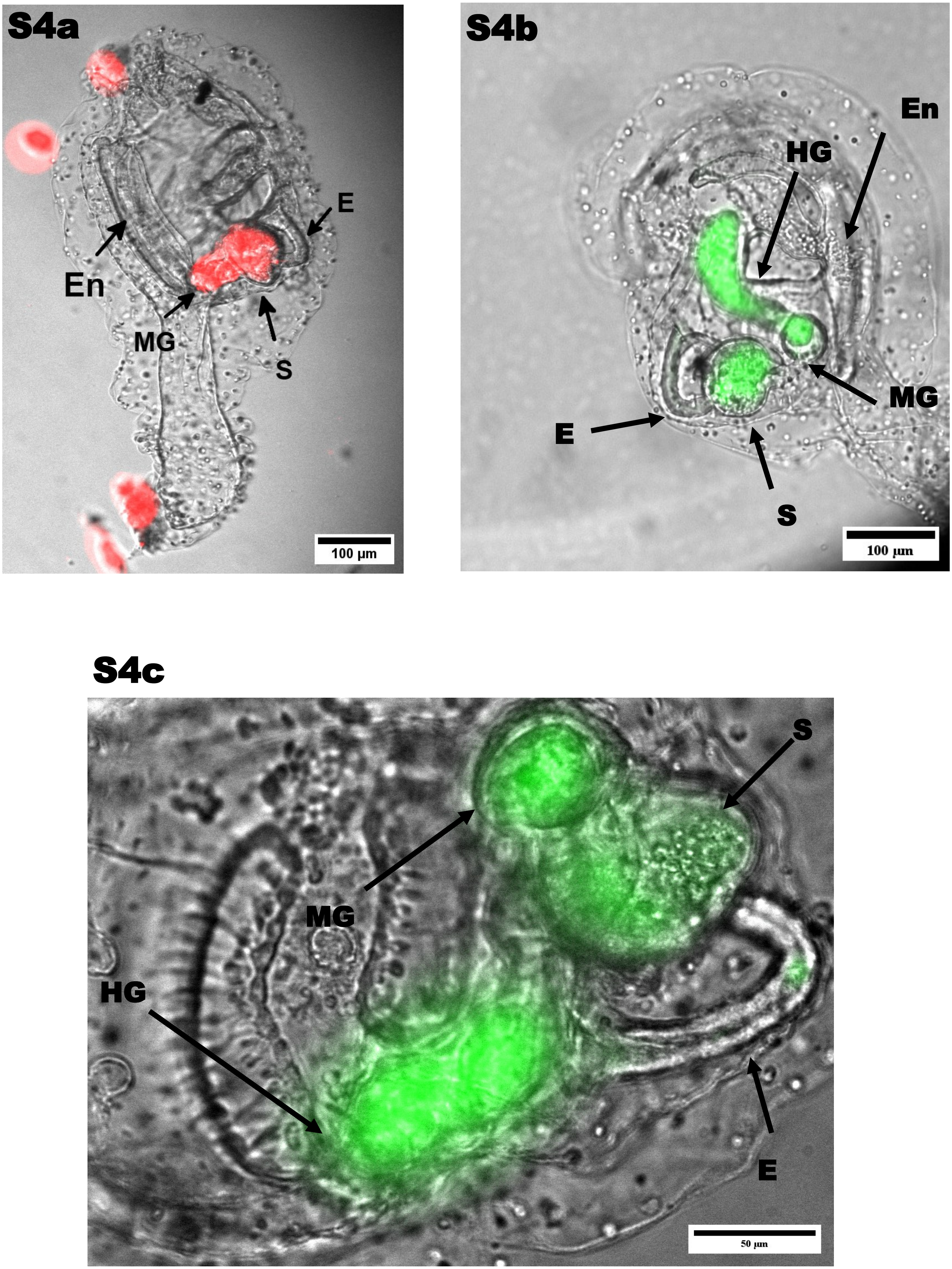
